# Massively parallel identification of sequence motifs triggering ribosome-associated mRNA quality control

**DOI:** 10.1101/2023.09.27.559793

**Authors:** Katharine Y. Chen, Heungwon Park, Arvind Rasi Subramaniam

**Affiliations:** Basic Sciences Division and Computational Biology Section of the Public Health Sciences Division, Fred Hutchinson Cancer Center, Seattle, WA 98109, USA; Molecular and Cellular Biology Program, University of Washington, Seattle, WA 98195, USA

## Abstract

Decay of mRNAs can be triggered by ribosome slowdown at stretches of rare codons or positively charged amino acids. However, the full diversity of sequences that trigger co-translational mRNA decay is poorly understood. To comprehensively identify sequence motifs that trigger mRNA decay, we use a massively parallel reporter assay to measure the effect of all possible combinations of codon pairs on mRNA levels in *S. cerevisiae*. In addition to known mRNA-destabilizing sequences, we identify several dipeptide repeats whose translation reduces mRNA levels. These include combinations of positively charged and bulky residues, as well as proline-glycine and proline-aspartate dipeptide repeats. Genetic deletion of the ribosome collision sensor Hel2 rescues the mRNA effects of these motifs, suggesting that they trigger ribosome slowdown and activate the ribosome-associated quality control (RQC) pathway. Deep mutational scanning of an mRNA-destabilizing dipeptide repeat reveals a complex interplay between the charge, bulkiness, and location of amino acid residues in conferring mRNA instability. Finally, we show that the mRNA effects of codon pairs are predictive of the effects of endogenous sequences. Our work highlights the complexity of sequence motifs driving co-translational mRNA decay in eukaryotes, and presents a high throughput approach to dissect their requirements at the codon level.

## Introduction

Translation and decay of mRNA are fundamental stages of gene expression whose interplay is crucial for determining steady-state protein levels in the cell. The protein coding region of mRNA has been recently recognized as an important determinant of mRNA stability^1–10^. Ribosome elongation rates can vary along the protein coding region, which is sensed by diverse regulatory factors to trigger mRNA decay^7–20^. Dysregulation of mRNA decay pathways has been linked to neurological diseases, autoinflammatory diseases, and cancer^21–26^.

Several motifs in the protein coding region of eukaryotic mRNAs have been associated with changes in mRNA stability^3,4,6,9–13,27,28^. Nonoptimal codons decrease ribosome elongation rates and trigger Not5-dependent mRNA deadenylation and decay^3,29–31^. Strong ribosome stalls caused by polybasic residues, poly-tryptophan sequences, and rare codon repeats trigger ribosome collisions and Hel2-dependent ribosome-associated mRNA quality control (henceforth RQC)^6,7,10,19,20,28,32,33^. Poly-proline sequences decrease ribosome elongation rate, but such slowdowns are thought to be resolved by eIF5A and not trigger mRNA quality control^34,35^. Ribosome profiling studies have identified several dipeptide and tripeptide motifs that are enriched at sites of ribosome stalls and collisions^36–39^. However, whether such motifs are sufficient to trigger mRNA quality control is not known. Ribosome stalling motifs in endogenous protein coding sequences often depend on a complex combination of amino acid residues in the nascent peptide^40–44^, and thus their relation to the simple repeat stalling sequences studied in reporter assays is not clear.

We recently developed a massively parallel reporter assay to identify coding sequence motifs triggering mRNA decay in human cells^27^. Using this assay, we found that translation of a diverse set of dipeptide repeats composed of bulky and positively charged amino acids are sufficient to trigger mRNA decay in human cells. Nevertheless, the molecular mechanism by which translation of these dipeptide repeats triggers mRNA decay in human cells remains unknown. Further, the extent to which translation of bulky and positively charged residues serves as an evolutionarily conserved signal for mRNA decay in other eukaryotes is unclear. Since co-translational mRNA decay pathways have been extensively studied in the budding yeast *S. cerevisiae*^7,45–48^, we sought to use this as an experimental model to dissect the molecular mechanism and sequence requirements of coding sequence-dependent mRNA decay. By extending our massively parallel reporter assay from human cells to *S. cerevisiae*, we identify several mRNA-destabilizing dipeptide motifs including combinations of bulky and positively charged residues, as well as proline-glycine and proline-aspartic acid dipeptide repeats. We define Hel2-dependent RQC as the major pathway regulating mRNA decay triggered by translation of these dipeptide repeats. Using deep mutational scanning, we further characterize the biochemical requirements at the codon level for bulky and positively charged dipeptide repeats in triggering Hel2-dependent mRNA decay. Together, our results highlight the diversity of coding sequence motifs triggering co-translational mRNA decay in *S*.*cerevisiae*, define the biochemical requirements for their mRNA-destabilizing effects, and reveal the extent of evolutionary conservation of these motifs across eukaryotes.

## Results

### A massively parallel reporter assay for mRNA effects in *S. cerevisiae*

To study the effect of coding sequence motifs on mRNA levels in *S. cerevisiae* in an unbiased manner, we modified a pooled reporter assay that we previously developed in mammalian cells^27^ (Fig. 1A). In our design for *S. cerevisiae*, a tandem 8× repeat of all possible codon pairs (4096 pairs in total) is inserted between the *PGK1* and *YFP* coding sequences. The 8× repetition amplifies the effect of each codon pair on mRNA levels. Each codon pair repeat is followed by a 24 nucleotide random barcode without stop codons, which enables their accurate quantification without sequence-dependent biases. Barcode sequences linked to each codon pair insert are identified by sequencing the plasmid library. We integrated the plasmid library into a noncoding region of chromosome I of *S. cerevisiae*, extracted mRNA and genomic DNA, and counted barcodes by high throughput amplicon sequencing. Barcode counts in the cDNA normalized by corresponding counts in the genomic DNA provide a relative measure of the steady-state mRNA level of each codon pair insert in our library. We further normalized mRNA levels by the median value across all inserts in the library to account for different sequencing depths and to facilitate comparison across experiments.

We recovered a median of 20 barcodes linked to each codon pair insert in the cDNA and genomic DNA libraries out of the 100 barcodes per insert in the plasmid library (Fig. S1A). We identified barcodes linked to 97% of all codon pairs in the plasmid library and 91% in the cDNA and genomic DNA libraries (Fig. 1B), indicating our assay’s ability to capture most of the codon pair motifs. Missing codon pairs in the plasmid library have a high GC content (Fig. S1B), suggesting that they are either resistant to cloning or toxic for *E. coli* growth. Many of the remaining missing codon pairs in the cDNA and genomic DNA from *S. cerevisiae* encode hydrophobic amino acids (Fig. S1C). Constitutive expression of such dipeptide repeats might be toxic due to their aggregation or membrane insertion.

**Fig 1:**
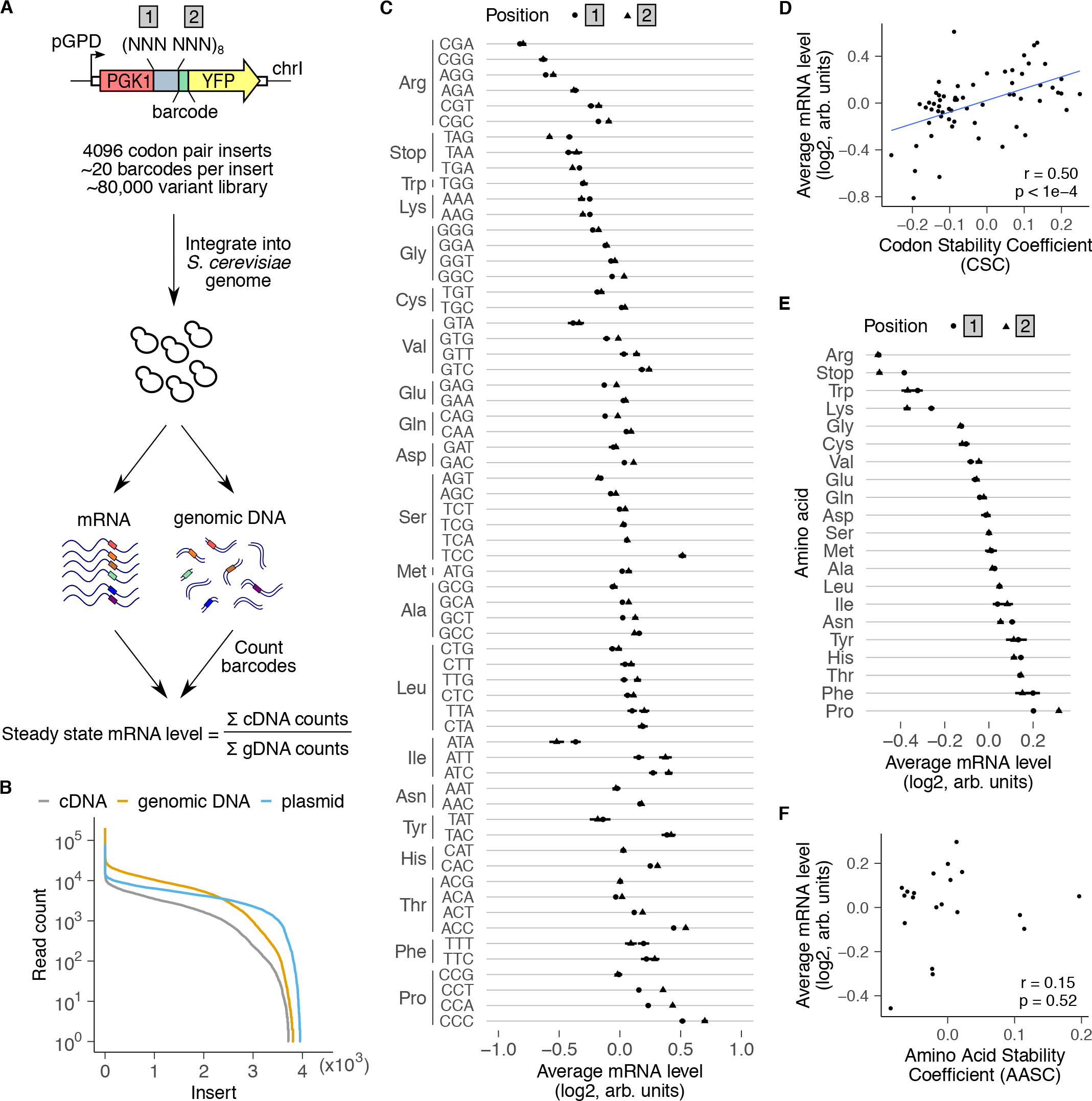
A massively parallel reporter assay for mRNA effects in *S. cerevisiae*. **(A)** Assay design. Each element in the library includes one of 4096 possible combinations of codon pairs repeated eight times. Each repeat is inserted in-frame between *PGK1* and *YFP*, and is followed by a random 24 nt barcode without in-frame stop codons (median of 20 barcodes/insert). The 80,000 variant library is integrated as a pool into a noncoding region of chromosome I. The barcodes in cDNA and genomic DNA are counted by high throughput amplicon sequencing. Relative steady state mRNA effect of each insert is calculated by first normalizing cDNA counts by genomic DNA counts for all barcodes linked to that insert and then by median-normalizing across all codon pairs. **(B)** Distribution of reads per codon pair insert for cDNA, genomic DNA, and plasmid libraries. **(C)** Average mRNA level of reporters with indicated codons in position 1 (circles) or position 2 (triangles) of the codon pair. **(D)** Average mRNA effects of individual codons compared against codon stability coefficients derived from endogenous *S. cerevisiae* mRNAs^3^. **(E)** Average mRNA level of reporters encoding the indicated amino acid in position 1 (circles) or position 2 (triangles) of the codon pair. Error bars in C and E represent standard deviation over all variants containing the codon or amino acid at each position. Average mRNA levels in C and E are median-normalized over all codons or amino acids at each position. **(F)** Same as D, except for amino acids compared against amino acid stability coefficients^4^.

To test whether our massively parallel assay recapitulates known codon and amino acid effects, we examined the average mRNA levels of individual codons and amino acids (Fig. 1C,E). To this end, we calculated the normalized ratio of barcode counts between cDNA and genomic DNA across all codon pairs containing each of the 64 codons or 20 amino acids. We observed a tight overlap of average mRNA levels of each codon or amino acid between positions 1 and 2 of the codon pair (Fig. 1C,E). This observation is consistent with the 8× repetitive nature of our codon pair library, due to which each codon pair insert is similar to its codon-reversed counterpart except for circular permutation of a single codon.

Within several synonymous codon families, codons with lowest mRNA levels in our assay (Fig. 1C) correspond to the less frequent codons within that family in the *S. cerevisiae* transcriptome^49–51^. These include CGA, CGG, and AGG (Arg), ATA (Ile), and CCG (Pro) (Fig. 1C), all of which are known to reduce protein expression or trigger mRNA decay in *S. cerevisiae*^3,5,6,20,52,53^. In line with these observations, average mRNA levels of codons in our assay positively correlated with codon stability coefficients (CSCs) inferred from stability measurements on endogenous mRNAs in *S. cerevisiae*^3,4^ (Fig. 1D, r=0.50, p<1e-4). This correlation with CSC is notable given that we vary only a 16 codon region within a 700 codon *PGK1-YFP* coding sequence in our assay.

At the amino acid level, arginine, lysine, and tryptophan had the lowest mRNA levels on average (Fig. 1E), consistent with the known role for these amino acids in triggering ribosome-associated quality control^6,7,12,20,28,46,54–57^. mRNA effects of these amino acids are comparable to that of stop codons, which trigger nonsense-mediated mRNA decay (NMD). In contrast to the codon effects, average mRNA levels of amino acids in our assay do not show significant correlation with amino acid stability coefficients (AASCs) inferred from stability measurements on endogenous mRNAs in *S. cerevisiae*^4^ (Fig. 1F). This lack of correlation is in line with the limited role of amino acid identity in determining global mRNA stability in *S. cerevisiae*^3,4^.

Overall, the average mRNA effects of codons and amino acids in our massively parallel reporter assay corroborate previously known stalling sequences in *S. cerevisiae* and show expected correlation with mRNA stability metrics inferred from endogenous mRNAs.

### Identification of codon pair repeats that reduce mRNA levels

Inclusion of all possible codon pair repeats in our library allowed us to next study the effect of pairwise codon and amino acid combinations on mRNA levels (Fig. 2A,B). We found a strong correlation (r=0.92, p<1e-10) between mRNA effects of codon pairs and their reverse counterparts, indicating the robustness of our measurements (Fig. S1D). We identified several families of synonymous codon pairs that consistently reduced mRNA levels relative to the remaining inserts in the library (black outlines, Fig. 2A,B). Among the most destabilizing codon families were those encoding lysine, arginine, and tryptophan repeats, in agreement with the average destabilizing effect of these amino acids (Fig. 1E).

**Fig 2:**
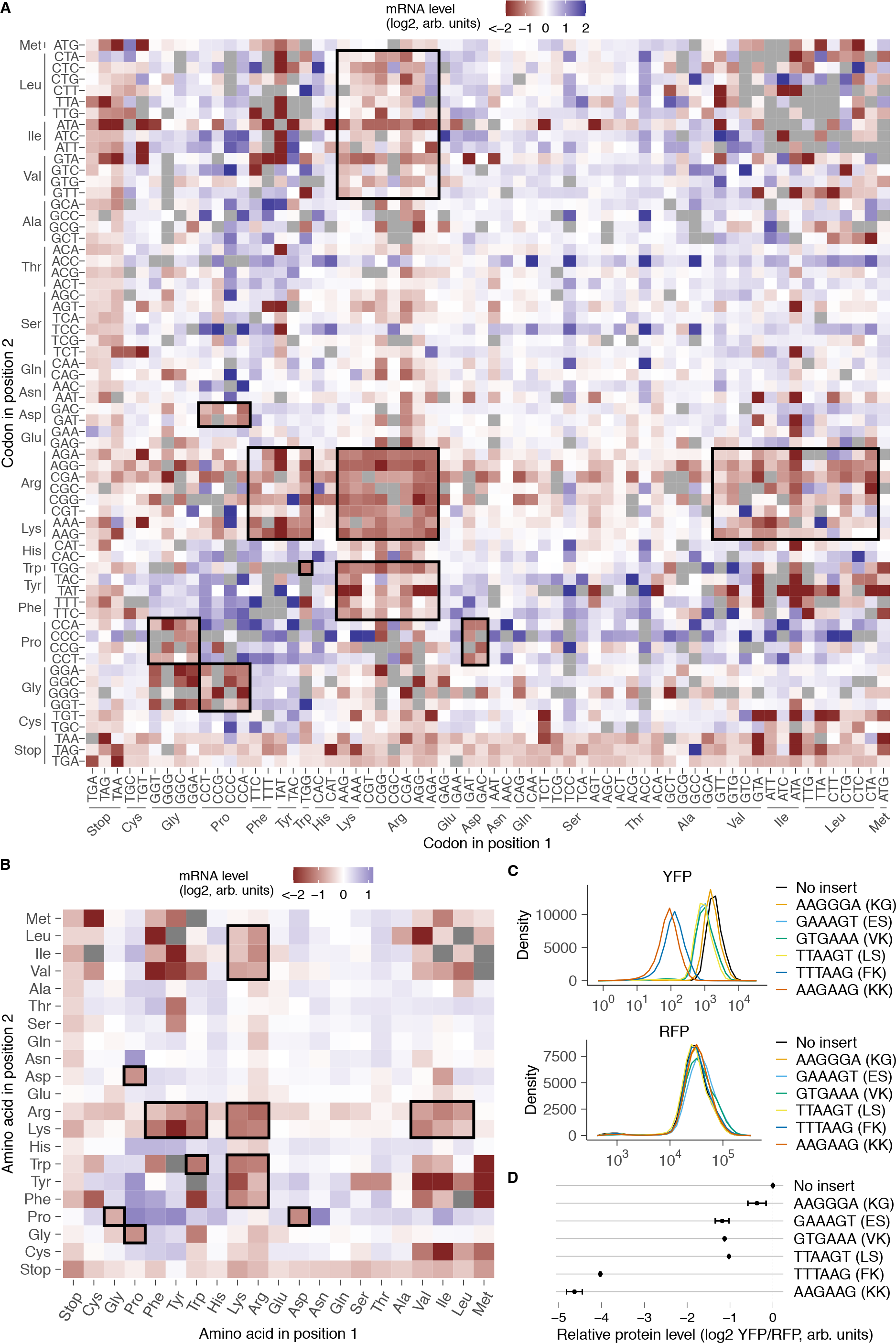
Identification of codon pairs and dipeptides that reduce mRNA levels. **(A)** mRNA level of inserts encoding each codon pair repeat. Codons at the first or second position of each pair are shown along the horizontal or vertical axes, respectively. Missing codon pairs are in grey. Synonymous codon pair families with lower mRNA levels are outlined in black. **(B)** mRNA level of inserts encoding each dipeptide repeat. Amino acids at the first or second position of each dipeptide are shown along the horizontal or vertical axes, respectively. Missing dipeptides are in grey. Dipeptide groups with lower mRNA levels are outlined in black. **(C)** Protein expression from individual *PGK1-YFP* reporters measured by flow cytometry (Top). A control RFP reporter integrated at a different locus was also quantified (Bottom). **(D)** Quantification of median YFP signal in **C** relative to the constitutively expressed RFP reporter. Error bars represent standard error of the mean across 5 biological replicates. GAAAGT (ES) is a frameshift control for GTGAAA (VK), and TTAAGT (LS) is a frameshift control for TTTAAG (FK).

Our assay revealed several dipeptide repeats that have not been previously associated with ribosome stalling or ribosome-associated quality control in *S. cerevisiae* (Fig. 2A,B). These include several combinations of bulky and positively charged amino acids such as phenylalanine-lysine (FK/KF), tryptophan-arginine (WR/RW), and tyrosine-lysine (YK/KY). Some combinations of hydrophobic and positively charged amino acids such as arginine-leucine (LR/RL) and arginine-isoleucine (IR/RI) were also destabilizing. Notably, we found similar mRNA-destabilizing combinations of positively charged amino acids with bulky and hydrophobic amino acids in human cells^27^, indicating that these sequences may be broadly destabilizing across eukaryotes. We confirmed the requirement of bulkiness for reducing mRNA levels in a targeted experiment by replacing phenylalanine with the smaller non-polar glycine in combination with lysine (Fig. S2A). Using flow cytometry, we found FK dipeptide repeats reduced YFP reporter levels similar to the known RQC-inducing KK repeat (Fig. 2C,D). Moreover, protein levels of a control RFP reporter expressed from a different chromosomal locus was unaffected by FK repeat expression, indicating that it does not perturb global gene expression (Fig. 2C).

Proline-glycine (PG/GP) and proline-aspartic acid (PD/DP) repeats were also among the mRNA-destabilizing codon pairs in our assay (black outlines, Fig. 2A,B). Unlike combinations of bulky and positively charged amino acids, these repeats did not reduce mRNA levels in human cells^27^. Conversely, amino acid combinations such as arginine-histidine and serine-phenylalanine that destabilize mRNAs in human cells^27^ did not reduce mRNA levels in our assay in *S. cerevisiae*. Finally, dipeptides comprised of bulky and positively charged amino acids as well as proline-glycine and proline-aspartic acid dipeptides are enriched at sites of ribosome collisions in *S. cerevisiae* and mammalian cells^36–38^. This observation suggests that the mRNA-destabilizing effects of such dipeptide repeats in our assay arises from ribosome slowdown when these peptide motifs are synthesized during mRNA translation.

### Dipeptide-induced mRNA destabilization requires translation

We used three different approaches to assay whether translation of dipeptide repeats is necessary for their mRNA-destabilizing effects.

First, we computationally tested whether the presence of codon pairs in the correct *PGK1-YFP* reading frame is necessary for the mRNA effects of the corresponding dipeptide repeats (Fig. 3A). mRNA effects of dipeptide repeats encoded in the correct +0 frame showed much lower correlation with the mRNA effects in the wrong +1 and +2 frames than with the correct +3 frame. We note that the +3 frameshift is essentially the same frame as the in-frame codon pair but with the codon positions interchanged. Thus, the simple presence of nucleotide sequences coding for destabilizing dipeptide repeats in the mRNA is not sufficient to reduce mRNA levels; they need to be present in the correct translated frame. Consistent with this observation, we found low correlation between mRNA levels of codon pair inserts and basic measures of nucleotide diversity such as GC content or GC3 content (Supplementary Fig. 2B).

**Fig 3:**
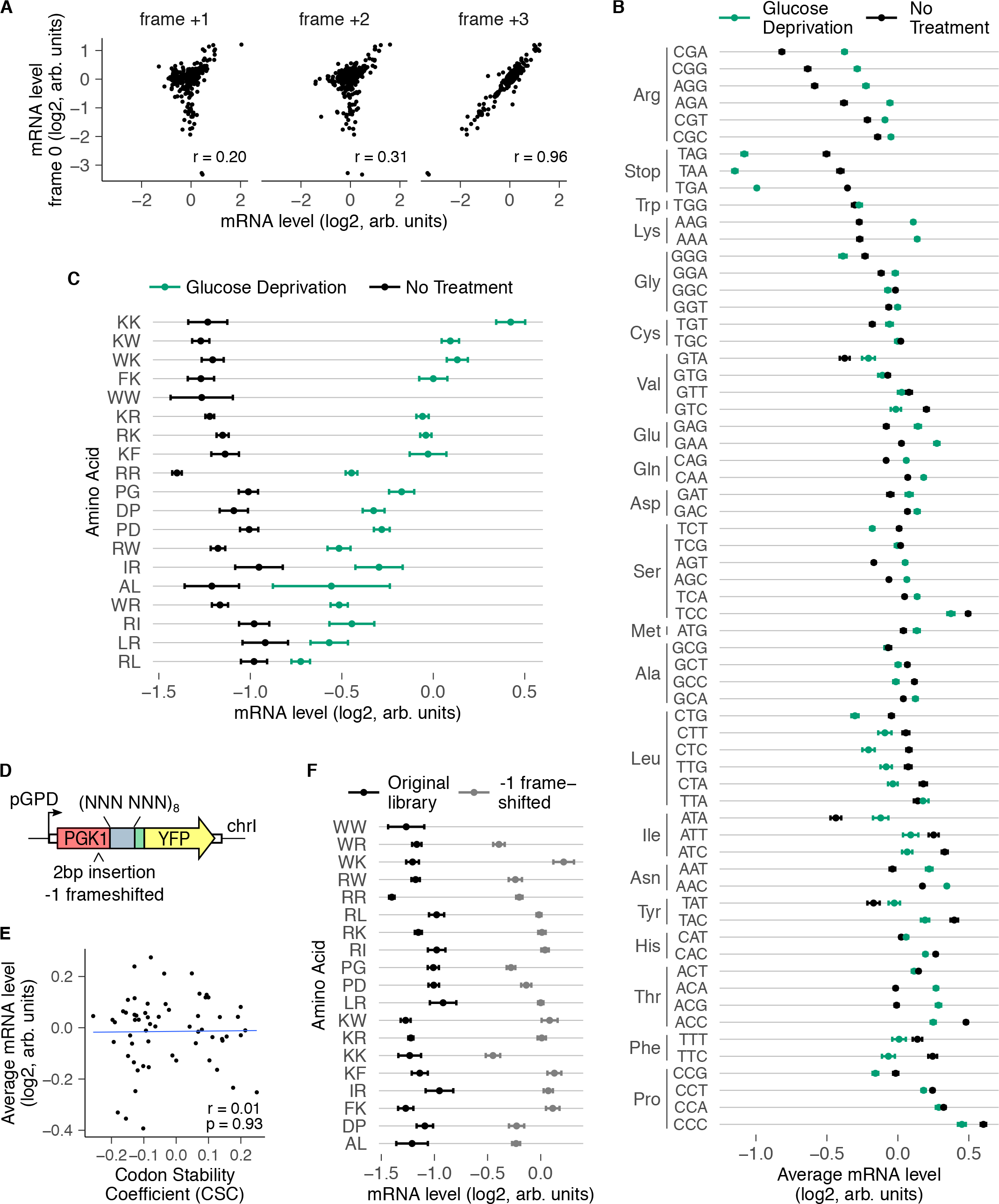
mRNA effects of dipeptide repeats require in-frame translation. **(A)** mRNA level of reporters encoding 320 different dipeptide repeats (excluding stop codon-containing dipeptides and pairs that did not pass read count cutoffs) compared between the correct reading frame (frame 0, vertical axis) and computationally-shifted +1, +2, or +3 reading frames (horizontal axes). *r* indicates Pearson correlation coefficient. **(B)** Average mRNA level of reporters with indicated codons averaged across positions 1 and 2 of the codon pair library during normal growth and glucose depletion. mRNA levels were median-normalized separately for each growth condition. Error bars represent standard deviation over all variants containing the codon at either position. **(C)** mRNA level of reporters encoding indicated dipeptides during normal growth and glucose deprivation. mRNA levels were median-normalized separately for each growth condition. Only dipeptide inserts with a minimum of 10 reads per barcode, 4 barcodes per insert, and low variability between barcodes are included here and in further analysis. Error bars represent standard deviation over barcodes linked to the indicated dipeptide repeat. **(D)** Schematic of frameshifted codon pair library. Two base pairs were inserted upstream of the codon pair insert in the 4096 codon pair library to create a -1 frameshift in the codon pair. Libraries were integrated and sequenced as in Fig. 1A. **(E)** Average mRNA effects of individual codons in the -1 frameshifted library compared against codon stability coefficients^3^. **(F)** mRNA levels of destabilizing dipeptides in the original in-frame library and in the -1 frameshifted library. Error bars calculated as in Fig. 3C.

Second, we tested whether global inhibition of translation is sufficient to rescue the mRNA-destabilizing effects of dipeptide repeats. Glucose deprivation is known to rapidly inhibit translation initiation in yeast^58,59^. Therefore, we grew *S. cerevisiae* cells containing the original codon pair library (Fig. 1A) in media without glucose for one hour, and quantified relative mRNA levels of inserts by high throughput sequencing as before. At the codon level, glucose deprivation increased the relative mRNA levels of inserts containing arginine and lysine codons, consistent with their mRNA effects arising at the translational level (Fig. 3B). Glucose deprivation also increased the relative mRNA levels of several dipeptide-encoding inserts that were destabilizing under normal growth (Fig. 3C). These include the known RQC-inducing polybasic sequences RR, RK, KR, and KK, as well as the novel destabilizing dipeptide repeats such as KW, FK, RW, PD, and PG that we identified in our original screen. Intriguingly, stop codon-containing inserts had lower mRNA levels during glucose deprivation even though nonsense-mediated mRNA decay of these inserts requires translation. This might be because NMD occurs in processing bodies (P-bodies), whose formation is enhanced upon glucose deprivation^60–62^.

Third, we tested whether experimentally altering the translated reading frame of codon pair inserts is sufficient to abrogate their mRNA-destabilizing effects, which would rule out transcription or RNA processing as possible mechanisms. Therefore, we inserted 2 base pairs upstream of the codon pair insert, leaving all other aspects of the reporter identical to the original library, and assayed for mRNA effects as before (Fig. 3D). The 2 base pair insertion shifts all codon pair inserts to the -1 frame, but does not introduce a stop codon upstream of the codon pair inserts. At the aggregate level, the -1 frameshifted library loses the previously observed correlation with codon stability coefficients (Fig. 3E, compare against Fig. 1D), consistent with the codon effects predominantly arising from translation. Similarly, most dipeptide repeats that destabilize mRNAs in the original library had higher relative mRNA levels in the -1 frameshifted library (Fig. 3F). Note that the WW dipeptide-coding repeat did not pass our read cutoff filter in both the glucose deprivation and the -1 frameshifting experiment (Fig. 3C,F).

In summary, our computational and experimental frameshifting assays, along with our glucose depletion experiment, establish the translation dependence of the mRNA effects of the destabilizing dipeptide repeats identified in our original screen.

### Ribosome-associated quality control regulates mRNA destabilization by dipeptide motifs

Given the translational dependence of mRNA destabilization by dipeptide repeats, we sought to identify the co-translational regulatory pathways mediating these effects. Ribosome stalling at poly-lysine, polyarginine, and poly-tryptophan repeats triggers ribosome-associated quality control (RQC) of nascent peptides and mRNAs in *S. cerevisiae*^6,28,55–57^. The E3 ubiquitin ligase Hel2 (*S. cerevisiae* homolog of human ZNF598), which binds collided ribosomes at extended ribosome stalls, is necessary for RQC induction at these sequences^10,32,56,57,63–65^ (Fig. 4A). Syh1 (GIGYF2 in humans) has also been recently implicated in a Hel2-independent pathway of mRNA decay of reporters with repeats of the rare codon CGA^66–68^ (Fig. 4A). To test the requirement for these factors in reducing the mRNA levels at the novel destabilizing dipeptide repeats identified in our screen, we integrated our original 4096-codon pair library into *S. cerevisiae* strains with *HEL2* or *SYH1* deletion, and measured relative mRNA levels as before (Fig. 4B).

**Fig 4:**
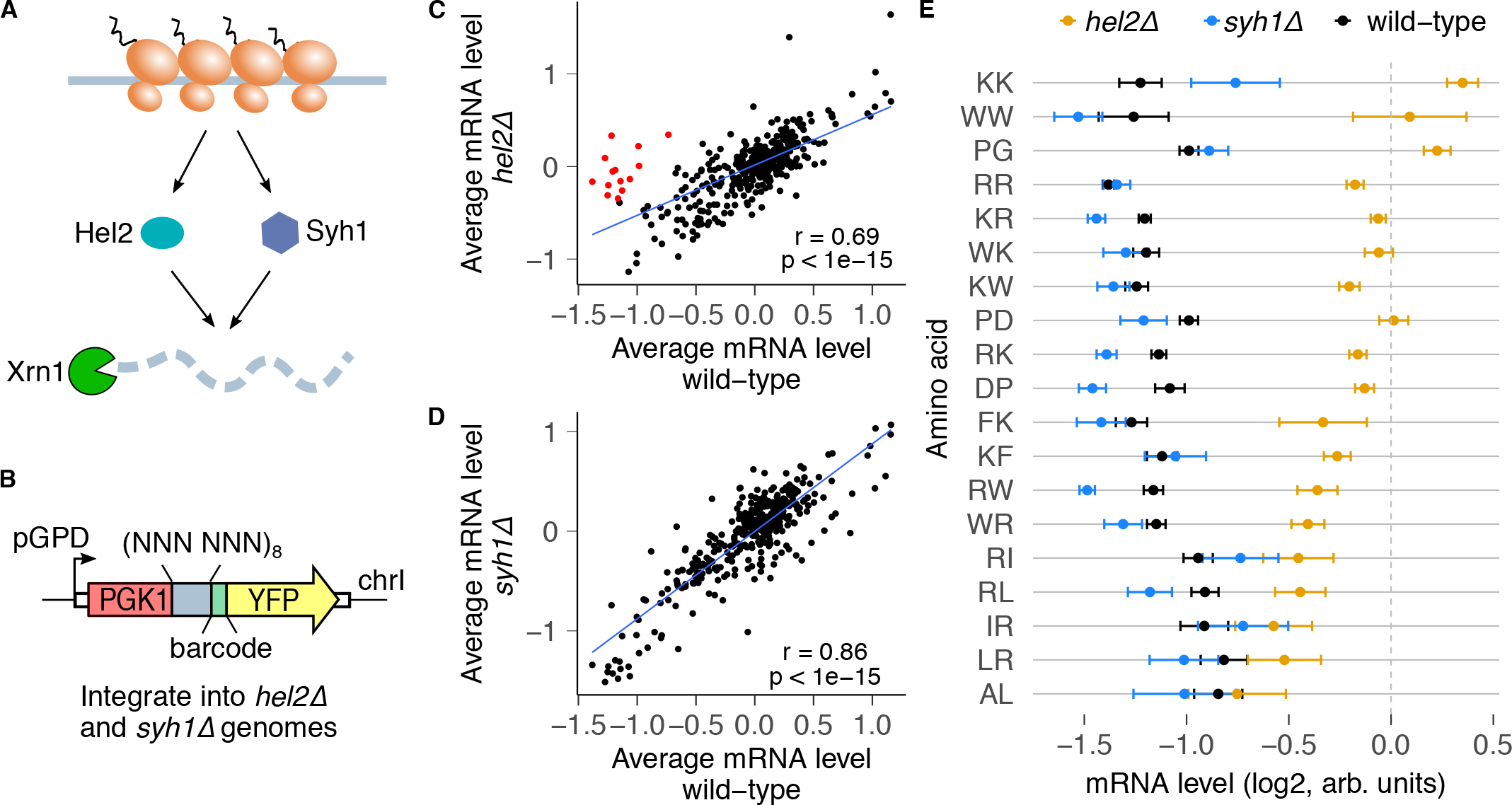
Ribosome collision sensor Hel2 regulates the mRNA effects of dipeptide repeats. **(A)** The RQC factors Hel2 and Syh1 are known to respond to collided ribosomes and trigger mRNA decay through Xrn1. **(B)** The codon pair library in Fig. 1A was integrated into *hel2Δ* and *syh1Δ* cells, and mRNA levels were quantified as before. **(C)** mRNA levels for dipeptide repeats compared between *hel2Δ* and wild-type cells. mRNA levels were calculated as in Fig. 3C, and median-normalized separately for each strain. Dipeptide repeats with residuals less than -2 from the linear regression line are marked in red. **(D)** Same plot as in C, but for *syh1Δ* cells. No dipeptide repeats are preferentially stabilized in *syh1Δ* cells with residuals less than -2 from the linear regression line. **(E)** mRNA levels for wild-type mRNA-destabilizing dipeptides (from Fig. 3C) for *hel2Δ* and *syh1Δ* cells. Error bars represent standard deviation over barcodes linked to the indicated dipeptide repeat.

We compared by linear regression the relative mRNA levels in the *hel2Δ* and *syh1Δ* strains against the wild-type strain to identify inserts with altered mRNA levels (Fig. 4C,D).

In the *hel2Δ* strain, 14 dipeptides had 1.5-fold or greater increase in relative mRNA levels compared to the wild-type strain (red points, Fig. 4C). These include the known RQC-inducing repeats, KK, RR, WW, RK, and KR. *HEL2* deletion also restored the mRNA levels of several bulky and positively charged dipeptide repeats (FK/KF, WR/RW, WK/KW) as well as proline-aspartic acid (PD/DP) and proline-glycine (PG/GP) repeats (Fig. 4E, Supplementary Fig. 3C). By contrast, *SYH1* deletion did not restore the mRNA levels of any dipeptide repeat (Fig. 4D,E). This is likely because Syh1 acts as a compensatory mechanism when Hel2-mediated RQC is inactive^66^. mRNA destabilization by a few combinations of positively charged and hydrophobic amino acids (RL/LR, RI/IR) was not rescued by either *HEL2* or *SYH1* deletion. Together, these results reveal that Hel2-mediated RQC regulates most but not all mRNA-destabilizing effects of dipeptide repeats identified in our original screen.

### Deep mutational scanning identifies critical residues mediating mRNA destabilization by dipeptide motifs

Ribosome-associated quality control often depends on interactions between specific residues in the nascent peptide and various regions of the ribosome such as the peptidyl-transferase center (PTC) and the uL4/uL22 constriction point in the exit tunnel^8,28,40,43,44^. To dissect the mechanism by which the FK dipeptide repeat triggers mRNA destabilization, we developed a deep mutational scanning assay using reporter mRNA level as a readout (Fig. 5A). Specifically, we mutated each location in the 16-codon insert encoding (FK)_8_ to all 64 codons to generate a pooled library of 1024 variants. We cloned these variants between the *PGK1* and *YFP* coding sequences, integrated them into the genomes of wild-type and *hel2Δ* cells, and measured variant frequency in cDNA and genomic DNA by high throughput amplicon sequencing. We used the ratio of cDNA to genomic DNA to quantify the relative mRNA levels of each variant, and further normalized to spike-in control strains to enable comparison across different genotypes (see Methods). We confirmed reproducibility of mRNA levels between biological replicate transformations into *S. cerevisiae* of the same plasmid library (Fig. 5B).

**Fig 5:**
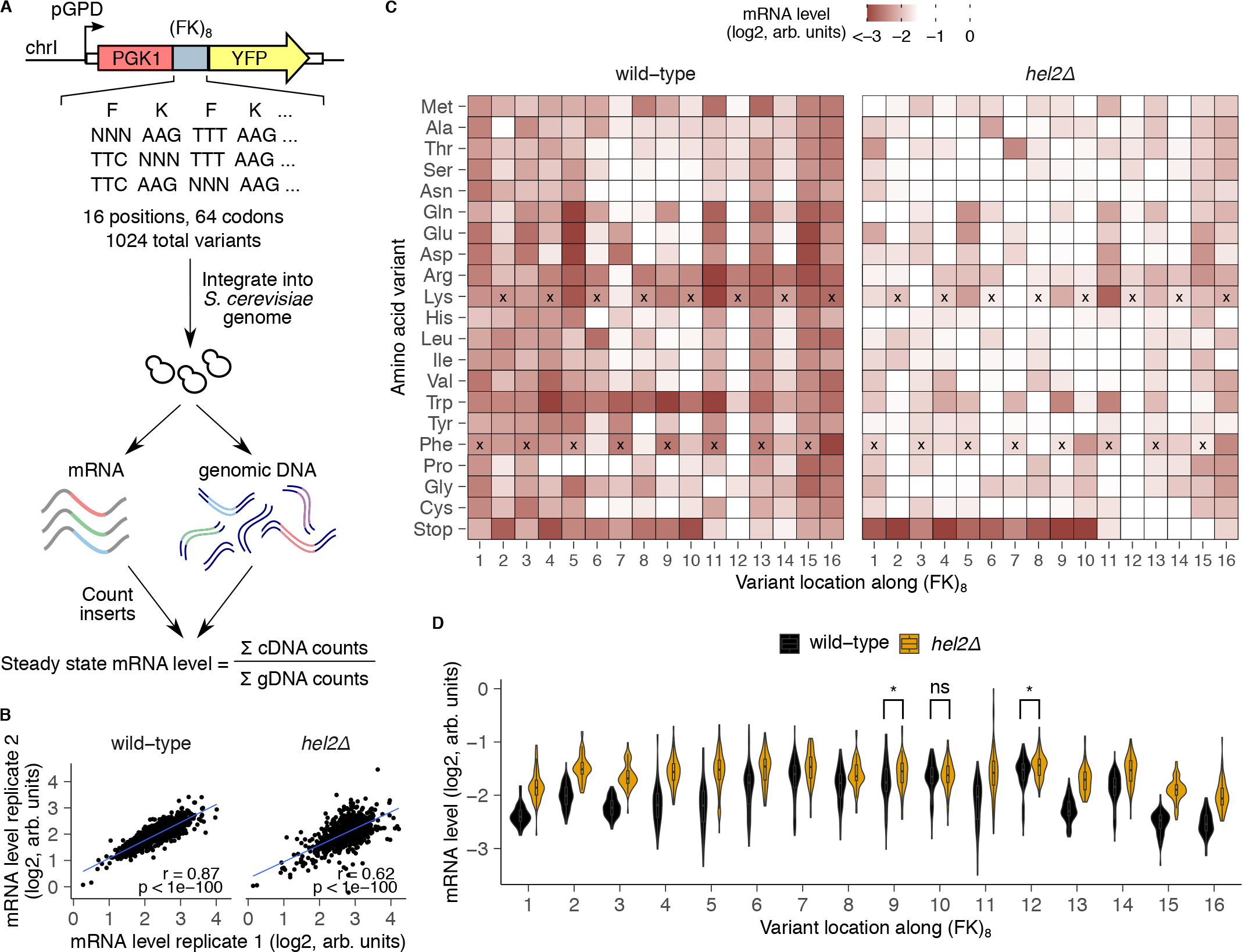
Deep mutational scanning identifies amino acids critical for mRNA effects of a destabilizing dipeptide repeat. **(A)** Schematic of deep mutational scanning (DMS) of the FK dipeptide repeat. Each location in an (FK)_8_-encoding insert was randomized to all 64 codons. This 1024-variant library was cloned as a pool between *PGK1* and *YFP*, and genomically integrated into wild-type and *hel2Δ* strains. Inserts were quantified in cDNA and genomic DNA by high throughput amplicon sequencing. **(B)** Pearson correlation between biological replicates for each variant in the (FK)_8_ DMS library. **(C)** mRNA level for inserts containing the indicated amino acid mutation (vertical axis) at the indicated position (horizontal axis). mRNA levels are averaged across replicates and normalized within each genotype using spike-in control strains. The wild-type amino acid variant is marked with black crosses at each location. **(D)** Violin plots of mRNA level across all amino acid variants at each location in wild-type and *hel2Δ* cells for both replicates combined. Stop codon variants are excluded from this analysis. Any locations where distributions were not significantly different (p>0.01 by Wilcoxon rank sum test) are marked.

Visualizing the relative mRNA levels of (FK)_8_ mutants as a function of mutation identity and location yields several interesting observations (Fig. 5C). First, reporter mRNA levels increase sharply when stop codons are present at positions 11 through 16 in wild-type cells. Since translation of premature stop codons will trigger mRNA decay through the Hel2-independent NMD pathway, our results imply that a minimum of 10 residues of the FK dipeptide need to be translated in order to trigger Hel2-driven RQC over NMD when the (FK)_8_ variants contain a stop codon. Interestingly, we also observe NMD suppression when stop codons are introduced after 10 residues of (FK)_8_ in *hel2△* cells, suggesting that extended ribosome stalling or collisions on mRNAs are sufficient to suppress NMD. Second, mRNA levels for nearly all mutations from positions 1 to 6 are as low as the wild-type sequence. This observation is again consistent with 10 residues in (FK)_8_ being the minimum RQC-inducing length, because mutating any of the first six residues will preserve this minimum length down-stream of the mutated position. Pro is the only target mutation within the first six positions that consistently rescues mRNA levels, likely by limiting the conformational flexibility of the nascent peptide^69–71^. Third, location 12 (and to a lesser extent location 14) within (FK)_8_ are the sole positions that require positively charged Arg or Lys to trigger Hel2-dependent RQC. At several other locations where the original amino acid is positively charged (such as at positions 6, 8, and 10), mutation to the bulkiest Trp residue can still trigger RQC, while mutations to other aromatic amino acids (Phe and Tyr) are insufficient. Fourth, at some locations where the original amino acid is bulky (such as at positions 9 and 11), mutating to the bulkier Trp or to positively charged Arg or Lys maintains RQC. The two preceding observations imply that positive charge and bulkiness play interchangeable roles at several locations within the (FK)_8_ repeat in triggering RQC. Finally, at position 7, where the original amino acid is Phe, mutations to other aromatic amino acids (Trp or Tyr) or to a negatively charged residue (Glu or Asp) triggers RQC, while positive charge is insufficient. Thus, the interchangeability of bulkiness with positive charge in triggering RQC is not universal, but rather depends on the location within the stalling peptide.

We next compared the aggregate effect of all mutations at each location of the (FK)_8_ repeat on mRNA levels between wild-type and *hel2△* cells (Fig. 5D). We excluded stop-codon containing mutants from this analysis to avoid convoluting NMD and RQC effects. The positions with the highest mutational effect differences between the two strains are at the ends of the stalling sequence: positions 1-6, 15, and 16 of (FK)_8_. This observation is consistent with our earlier interpretation that translation of 10 residues of (FK)_8_ is necessary to drive Hel2-dependent mRNA decay. Conversely, positions 10, 9, and 12 had the least mutational effect differences between the two strains, revealing that these positions are most important for triggering Hel2-dependent RQC. Finally, *HEL2* deletion did not fully rescue the mRNA effects of any (FK)_8_ terminal mutants (positions 1, 15, 16), suggesting that Hel2-dependent RQC activity is saturated at longer repeat lengths, and mRNA decay proceeds through multiple compensatory pathways.

### Codon pair library predicts mRNA effects of endogenous sequences

Though a few mRNA sequences are known to stall ribosomes and trigger RQC in reporter studies^40,41,72^, the sequence motifs that underpin endogenous mRNA stability are not well understood. For example, the simple presence of polybasic stretches or rare codons is not sufficient to trigger quality control on endogenous yeast mRNAs^40,73^. Thus, we sought to test whether our codon pair assay could predict mRNA effects of sequence motifs in endogenous *S. cerevisiae* genes. To this end, we assayed 1904 fragments, each 48 nucleotides long, from endogenous mRNAs spanning a wide range of expression levels^74^ using the same reporter design as the codon pair library (Fig. 6A). We integrated this endogenous fragment library into wild-type cells and counted barcodes by high throughput amplicon sequencing as before. Compared to the codon pair library, mRNA levels in the endogenous fragment library were more tightly distributed around the median, indicating more muted effects on mRNA stability (Fig. 6B). We next calculated the codon stability coefficient (CSC) values for each of the 64 codons using mRNA levels either from the codon pair library or the endogenous fragment library^3^. We found strong correlation (r=0.67, p<1e-8) between the two libraries, indicating that mRNA effects of codon pair repeats predict mRNA effects of endogenous sequence motifs in wild-type cells (Fig. 6C). We next integrated the endogenous fragment library into *hel2△* cells and tested how Hel2-dependent RQC affects the relationship between CSC values calculated from the codon pair and the endogenous fragment libraries. We found that *hel2△* cells still exhibited a significant correlation (r=0.49, p<1e-4) between the two libraries, though to a lesser extent than in wild-type cells (Fig. 6D), consistent with the majority of endogenous sequences not triggering prolonged ribosome slowdown or collisions.

**Fig 6:**
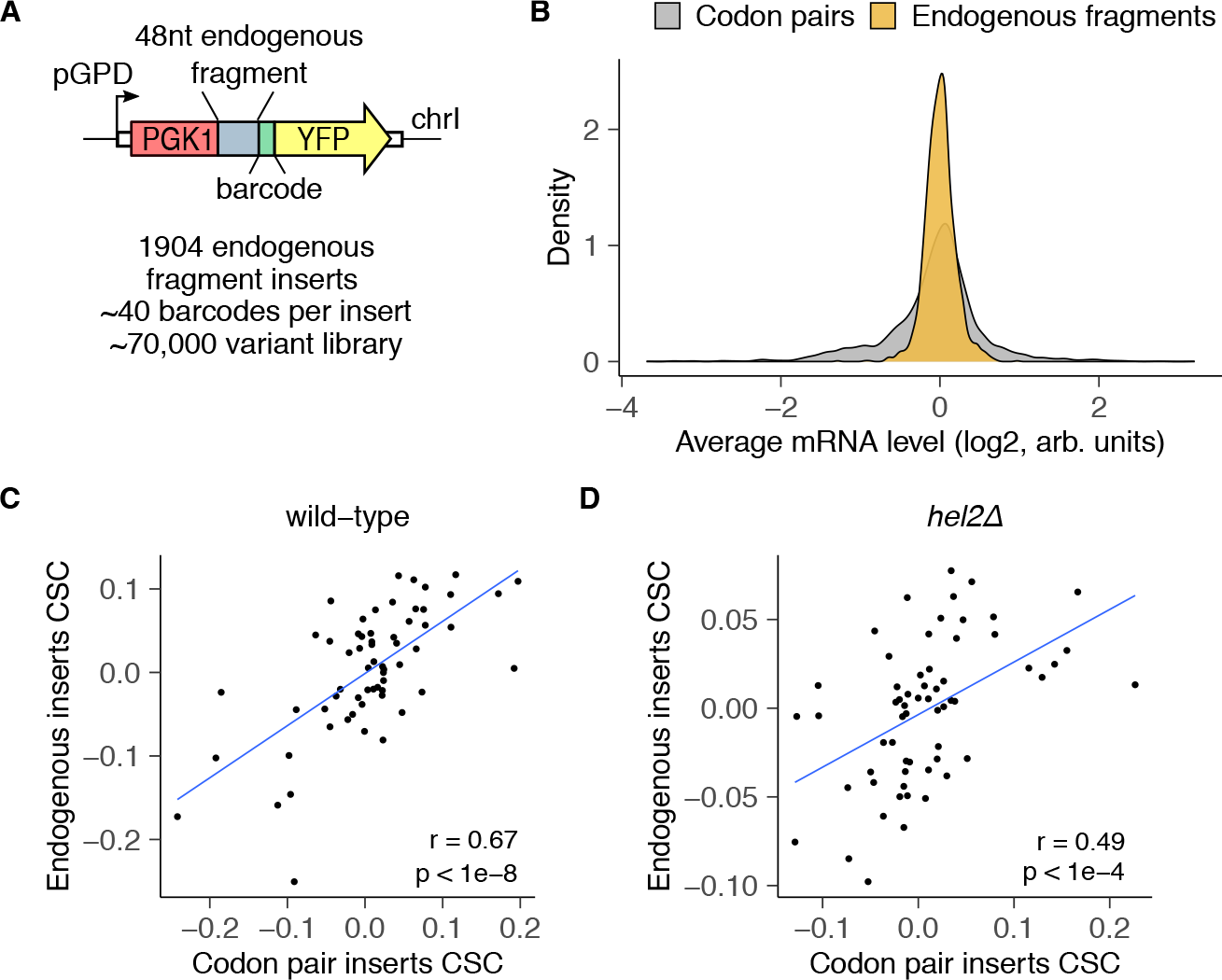
Codon pair measurements predict effects of endogenous mRNA sequences. **(A)** Schematic of endogenous sequence insert library. Each element in the library includes one of 1904 possible 48nt endogenous fragments. Each sequence is inserted in-frame between *PGK1* and *YFP*, and is followed by a random 24nt barcode without in-frame stop codons (median of 40 barcodes/insert). The 70,000 variant library is genomically integrated into wild-type and *hel2△* cells, and mRNA levels are quantified as in Fig. 1A. **(B)** Distribution of mRNA levels for endogenous fragments vs codon pair inserts in wild-type cells. **(C)** Correlation between CSC values calculated for each codon from the endogenous fragment library against CSC values derived from the codon pair library in wild-type cells. Pearson correlation coefficient is reported as *r*. The CSC for each codon is calculated by taking the Pearson correlation coefficient between codon frequency of an insert and its steady state mRNA level. **(D)** Same plot as in C, but for *hel2△* cells.

## Discussion

Here, we use a massively parallel approach to identify and dissect sequence motifs underlying mRNA instability in *S. cerevisiae*. In addition to validating known codon and amino acid effects on mRNA stability, we identify several sequence motifs that have not been previously associated with mRNA decay. These include combinations of bulky and positively charged amino acids, and proline with asparate and glycine, all of which trigger translation-dependent mRNA decay through the Hel2-dependent RQC pathway. By combining our massively parallel assay with deep mutational scanning, we dissect the codon-level biochemical requirements for triggering mRNA decay by a bulky and positively charged dipeptide repeat. Despite the apparent simplicity of the codon pair repeat library, we find that it captures the mRNA effects of endogenous coding sequence fragments from the *S. cerevisiae* transcriptome.

Our codon pair library confirms the role of codon optimality as a major determinant of mRNA stability in *S. cerevisiae*, and provides insights into the resulting hierarchy of effects. We observe several synonymous codon families within which aggregate mRNA levels differ based on the hierarchy of codon optimality^3,52^ (Fig. 1C), but have different absolute effects. The nonoptimal codons ATA (Ile), GTA (Val), and TAT (Tyr) are highly destabilized relative to their optimal counterparts. By contrast, the optimal codon TCC (Ser) is preferentially stabilzed relative to its nonoptimal counterparts. Both the arginine and proline synonymous codon families are stratified based on codon optimality even though these amino acids have opposite average effects on mRNA stability (Fig. 1E, Arg – destabilizing, Pro – stabilizing). Thus, codon optimality effects on mRNA stability act in parallel and independent of amino acid identity. Consistent with codon optimality-mediated mRNA decay being a cotranslational process^29,75,76^, translational shutoff by glucose depletion rescues the mRNA-destabilizing effects of eight out of the 10 most non-optimal codons (ATA, CGA, AGG, GTA, ACG, AGT, AAA, AGC)^3^ (Fig. 3B). Finally, the effects of codon optimality on mRNA stability in our codon pair library are driven by mutations within a short 16 codon region despite being part of a 700 codon *PGK1-YFP* mRNA. This is likely because the *PGK1-YFP* region is efficiently translated^77^, while the tandem and repetitive nature of the codon pairs amplifies their effect on ribosome slowdown and recruitment of mRNA-destabilizing factors.

While polybasic and poly-tryptophan sequences are known to trigger RQC in *S. cerevisiae*, our codon pair assay reveals combinations of bulky (Val, Ile, Leu, Phe, Tyr, Trp) and positively charged (Arg, Lys) amino acids as a general trigger of mRNA decay (Fig. 2A,B). Interestingly, combinations of Val, Ile, Leu, and Phe with Arg and Lys were also found to destabilize mRNA in human cells^27^, indicating their evolutionary conservation as mRNA-destabilizing sequences across eukaryotes. Supporting these findings, ribosome profiling in human cells revealed an enrichment in disome occupancy at sites that followed an Arg-X-Lys pattern, with highest disome density occurring when X was Phe, Ile, or Leu^36^. We find that positively charged amino acids in combination with the bulkiest side chains (Phe, Trp) trigger RQC-dependent mRNA decay in *S. cerevisiae*, while less bulky side chains (Val, Ile, Leu) decrease mRNA levels in a Hel2-independent manner (Fig. 4E). We speculate that such mRNA motifs that stall ribosomes sufficiently to trigger mRNA decay but are less terminally stalling than Phe/Trp in combination with Arg/Lys may be acted on by compensatory pathways such as Syh1/Smy2^66^, or through Not5-dependent decay^29^.

In our codon pair assay, combinations of proline with aspartate and glycine (PD/DP, PG/GP) decrease mRNA levels in a Hel2-dependent manner (Fig. 2A,B, Fig. 4E, Supplementary Fig. 3). While poly-proline sequences stall ribosomes due to inefficient peptide bond formation, these sequences are not known to induce RQC and are instead translated with the assistance of eIF5A^34^. Consistent with these previous findings, proline-proline combinations, and all other proline-containing combinations except for with aspartate and glycine, are stabilizing in our assay. Conversely, no other aspartate or glycine containing codon pairs except the ones with proline are destabilizing. While increased ribosome occupancy has been observed at proline, aspartate, and glycine codons in both *S. cerevisiae* and human cells^36,78,79^, our results suggest that these effects may be driven by combinations of these amino acids rather than by their individual occurrence. Consistent with this idea, PD and PPD peptides have increased ribosome occupancy and are under-represented in the *S. cerevisiae* proteome, while PP and GG dipeptides also have increased ribosome occupancy but are over-represented^37^. Similarly, PD dipeptides in *E. coli*^80^, and PD and PG motifs in mouse embryonic stem cells^38^ have increased ribosome occupancy. Thus, PD and PG motifs may have evolutionarily conserved effects on ribosome slowdown through a mechanism distinct from poly-proline stalls, and can trigger Hel2-dependent mRNA decay in *S. cerevisiae*.

Our deep mutational scanning reveals complex codon-level requirements for the (FK)_8_ repeat to confer mRNA instability in a Hel2-dependent manner (Fig. 5). Strikingly, these results also exhibit several similarities to the composite biochemical requirements for ribosome stalling observed at the known endogenous RQC substrate in *S. cerevisiae, SDD1*_196-212_ (FFYEDYLIFDCRAKRRK)^40^. First, the strict requirement for positive charge at positions 12 and 14 of the (FK)_8_ repeat to trigger mRNA decay matches the requirement for positive charge at positions 207 and 209 of *SDD1*_196-212_, which are thought to perturb the peptidyl-transferase center of the ribosome. Second, the requirement for bulky aromatic residues at position 7 of (FK)_8_ is similar to the requirement for aromatic residues at position 201 of *SDD1*_196-212_, which are thought to interact with the uL4/uL22 constriction point of the ribosome. Third, the ability of negatively charged aspartate, and to a lesser extent glutamate, at position 7 of (FK)_8_ to preserve stalling resembles the requirement for aspartate at position 200 of *SDD1*_196-212_, though in the *SDD1* case, the requirement for aspartate is strict. Our results show that bulkiness can be compensated by negative or positive charge in stall sequences depending on the position along the sequence. Specifically, aspartate’s prevalence in stalling sequences is evident in ribosome profiling studies from *S. cerevisiae* to humans, which show increases in monosome and disome occupancy at aspartate codons^36,78,79^, presumably due to interactions with the negatively charged ribosome exit tunnel. Taken together, our deep mutational scanning results with a simple (FK)_8_ repeat recapitulate and generalize the biochemical requirements for ribosome stalling and quality control observed with endogenous stall sequences.

While we did not intend to focus on NMD for this study, our assay nonetheless identified several patterns related to NMD. Surprisingly, we found that glucose depletion selectively destabilized stop codon-containing mRNAs for all three stop codons (Fig. 3B) even though NMD depends on mRNA translation. A possible basis for this observation is that glucose depletion increases the formation of P-bodies, which sequester translationally silenced mRNAs including those subject to NMD^60,61^. Thus, the co-localization of NMD substrates with NMD factors at P-bodies might enhance their decay during glucose depletion^60,62,81,82^. Deep mutational scanning of the (FK)_8_ dipeptide also revealed the differential kinetics between NMD and RQC when in competition for the same substrates (Fig. 5C). Before 10 Phe and Lys residues are translated, stop-codon containing sequences are predominantly degraded by NMD. After this minimum stalling sequence is translated, RQC dominates as the primary regulatory mechanism. A minimum length of 10 Phe and Lys residues of RQC is consistent with 12 repeated tryptophan residues being sufficient to induce RQC, while greater than 8 residues were required^28^. Interestingly, in *hel2△* cells we observe that NMD is suppressed after (FK)_5_ repeats are translated, even though Hel2-dependent RQC and NMD should presumably not be competing in these cells. This suggests that extended ribosome stalling and collisions is sufficient to prevent degradation of NMD substrates.

The results of our combinatorial codon pair and endogenous motif mRNA stability assays suggest that a wider diversity of mRNA sequences impact mRNA stability than previously appreciated. Poly-GP repeats, identified in our study to stall ribosomes and trigger RQC, are translated through repeat associated non-ATG (RAN) translation of the pathogenic G_4_C_2_ repeat expansion in the *C9ORF72* gene and is a biomarker for C9ORF72-associated ALS^83^. Valine-arginine repeats, identified in our study to destabilize mRNAs in a Hel2-independent manner, are also translated through RAN in the mammalian TERRA sequence to form inclusions during disrupted telomere homeostasis^84^. Thus the sequences identified in our study have important implications in the maintenance of cellular homeostasis and disease progression.

## Author Contributions

K.Y.C. conceived the project, designed research, performed experiments, analyzed data, and wrote the manuscript. H.P. designed research and performed experiments. A.R.S. conceived the project, designed research, analyzed data, wrote the manuscript, supervised the project, and acquired funding.

## Acknowledgements

We thank Phil Burke, members of the Subramaniam lab and the Zid lab for discussions and feedback on the manuscript. This research was funded by NIH R35 GM119835 and NSF MCB 1846521 received by ARS. This research was supported by the Genomics Shared Resources of the Fred Hutch/University of Washington Cancer Consortium (P30 CA015704) and Fred Hutch Scientific Computing (NIH grants S10-OD-020069 and S10-OD-028685). The funders had no role in study design, data collection and analysis, decision to publish, or preparation of the manuscript.

## Data and Code Availability

The raw sequencing data generated in this study have been deposited in the Sequence Read Archive under BioProject accession number PRJNA974090, at https://www.ncbi.nlm.nih.gov/bioproject/?term=PRJNA974090. Raw data from flow cytometry are available at http://flowrepository.org/id/FR-FCM-Z6QH. Code to reproduce figures in the manuscript starting from raw data is publicly available at https://doi.org/10.5281/zenodo.8365102 and https://github.com/rasilab/chen_2023. Software environments used to run the code in the above GitHub repository are publicly available as Docker containers at https://github.com/orgs/rasilab/packages. Biological reagents or methodology clarification can be publicly requested by opening an issue at https://github.com/rasilab/chen_2023/issues.

## Materials and Methods

### Parent vector construction

Plasmids constructed and used in this study are listed in table S1. Oligonucleotides used in this study are listed in table S3. Plasmid assembly was carried out using standard molecular biology techniques as described below. All polymerase chain reaction (PCR) reactions were performed using Phusion polymerase (Thermo Fisher F530S) or Phusion Flash High-Fidelity PCR Master Mix (Thermo Fisher F548L) according to manufacturer’s instructions. Restriction enzymes were obtained from Thermo Fisher and FastDigest (FD) variants were used when available.

The chrI-integrating parent vector pHPSC1120 used for this study was constructed from pHPSC417 used in our previous work^20^. In comparison to pHPSC417, pHPSC1120 contains an additional Illumina Read1 primer binding site and T7 promoter sequences for deep sequencing of inserts and barcode sequences and for in vitro transcription from genomic DNA, respectively. The Illumina R1 sequencing primer binding and T7 promoter sequences were PCR-amplified using oHP558 as the forward primer, oHP530 as a bridge primer, and oHP529 as a reverse primer, and cloned into BamHI-linearized pHPSC417 using Gibson assembly. The -1 frameshifted parent vector pHPSC1114 was also constructed from pHPSC417 using the same strategy as for pHPHS1120 but with a different forward primer oHP528 that incorporates the frameshift. All plasmids were verified by Sanger sequencing.

### Variable oligo pool design

#### Pool 1

Pool 1 includes the 8× dicodon library (4096 codon pair inserts) and the endogenous gene fragments library (1904 inserts). The 8× dicodon library (Fig. 1A) encodes all possible codon pair (6 nucleotide) combinations, for a total of 4096 codon pairs. Each codon pair is repeated eight times to create 48 nucleotide (nt) inserts. The endogenous gene fragments library includes 1904 endogenous fragments, each 48 nt in length (Fig. 6A). Endogenous gene fragments were selected as 253 nt to 300 nt of each ORF. Only ORFs designated as “Verified” by the Saccharomyces Genome Database (SGD) in the R64-1-1 release were included (http://sgd-archive.yeastgenome.org/sequence/S288C_reference/genome_releases/). Every 2nd gene in descending order of RNA expression^74^ was included in this library to encompass a wide range of expression levels. All 6000 inserts are flanked with the same 29 nt 5’ homology arm and 24 nt3’ homology arm. The oligo pool (oAS385) was ordered from Twist Biotechnologies.

#### Pool 2

The FK_8_ deep mutational scanning library (Fig. 5A) was constructed from a starting sequence composed of phenylalanine and lysine codons repeated eight times in tandem (48 nt inserts). The phenylalanine codons TTT and TTC and the lysine codons AAA and AAG were used interchangeably throughout the insert to avoid producing a repetitive mRNA sequence. At each of the 16 positions, an NNN sequence was used to randomize the codon. The oligo pool (oKC224) was ordered as an oPool from Integrated DNA Technologies.

### Plasmid library construction

For the 8× dicodon library, oligo pool 1 (described above) was PCR-amplified with oKC97 and oHP531. For the -1 frameshifted 8× dicodon library, pool 1 was PCR-amplified with oHP532 and oHP531. As described above, oHP531 encodes a 24 nt random barcode region, comprised of 8× VNN repeats. Barcoded oligo pools were cloned into BamHI-linearized pHPSC1120 and pHPSC1114 by Gibson assembly. Assembled plasmid pools were transformed at high efficiency into NEB 10-Beta *E. coli* cells, and plated as 1:10 serial dilutions. 500,000 colonies were scraped from plates for extraction in order to bottleneck the number of unique variants. Pool 2 was PCR-amplified with oKC97 and oKC225 and cloned into BamHI-linearized pHPSC1120 by Gibson assembly. The assembled plasmid pool was transformed at high efficiency into NEB 10-Beta *E. coli* cells. 70,000 colonies were scraped from plates for extraction in order to bottleneck the number of unique variants.

### Individual plasmid construction

To generate the *PGK1-YFP* reporters used for flow cytometry of individually selected codon pairs, the desired codon pair inserts were amplified using two rounds of PCR from a pooled plasmid template pHPSC1136 not used in this study. Unique primers (oKC129-142) were used to amplify the six desired inserts. Homology arms were added to the six amplified inserts using oKC97 and oKC123 primers. Amplified inserts were cloned into BamHI-linearized pHPSC1120 by Gibson assembly to produce pHPSC1144, pHPSC1145, pHPSC1146, pHPSC1147, pHPSC1149, pHPSC1150 plasmids. All individual plasmids were verified by Sanger sequencing.

To create the small barcoded pool for mRNA measurement validation (Fig. S2A,E), oKC97 and oKC148 oligos were used to barcode and amplify inserts from the following plasmids (described above): pHPSC1144, pHPSC1145, pHPSC1146, pHPSC1147, pHPSC1149, pH-PSC1150. oKC148 encodes a 24 nt random barcode region, comprised of 8× VNN repeats. Barcoded inserts were then pooled at equimolar concentrations and cloned into BamHI-linearized pHPSC1120 by Gibson assembly. The assembled plasmid pool was transformed at high efficiency into NEB 10-Beta *E. coli* cells. 2,000 colonies were scraped from plates for extraction in order to bottleneck the number of unique variants. Two colonies were picked and Sanger sequenced to obtain the identity of the insert and barcode pair of the two spike-in plasmids, pHPSC1159-sc2 and pHPSC1159-sc5.

### Strain construction

All *S. cerevisiae* strains used in this study are listed in table S2. Integration of pooled plasmids into the *S. cerevisiae* genome was performed by transforming 30–200 μg of NotI-linearized plasmid library into 1– 5e9 cells according to the LiAc/SS carrier DNA/PEG method^85^. Following heat shock, cells were transferred into a 5x volume of a 1:1 solution of 20% dextrose and synthetic complete (SC) media lacking uracil with 2% dextrose (SCD-URA) and spun at 1850*g* for 5 minutes. Cell pellets were gently resuspended in 100mL of fresh SCD-URA and allowed to recover overnight at 30°C shaking at 200rpm. After 20–24 hours, 1e9 cells were passaged into 100mL fresh SCD-URA and grown overnight at 30°C shaking at 200rpm. This process was repeated for a total of 72 hours of selection in SCD-URA before making glycerol stocks from saturated cultures. Integration of individual constructs into the *S. cerevisiae* genome was performed by transforming 0.5– 1.0μg of linearized plasmid according to the LiAc/SS carrier DNA/PEG method^85^. Single yeast colonies were selected on SCD agar plates lacking uracil after 48 to 72 hours growth at 30°C.

### Harvesting pooled library cells

Glycerol stocks of cells containing pooled reporter strains were thawed and grown overnight in 20-50mL YEPD at starting OD_600_ between 0.1 and 0.5 at 30°C with shaking at 200rpm. The saturated cultures were diluted approximately 200-fold (for starting OD_600_ of 0.1) and spike-in strains (scKC190 and scKC191) were introduced into each culture at a concentration approximately the same as each library variant based on OD_600_ density. Cultures were grown for 4–6 hours at 30°C with shaking at 200rpm until mid-log phase (OD_600_ between 0.4-0.6), then transferred to ice-water baths. Each culture was split into 50mL aliquots (approximately >=200 million cells) in pre-chilled conical tubes and spun down at 3000*g*, 4°C, for 5 minutes. The supernatant was removed and the cell pellets were flash-frozen in a dry ice-ethanol bath and stored at -80°C.

### Harvesting glucose-depleted cells

Glycerol stocks of cells containing the pHPSC1142 pooled reporter library were thawed and grown overnight as described above. Saturated cultures were diluted and spike-in strains (scKC190 and scKC191) were introduced as described above. Cells were grown for 4 hours at 30°C with shaking at 200rpm until OD_600_ of 0.4. Cells were spun down at 3000rpm for 2 minutes and washed with 30mL H_2_O twice, then resuspended into YEP (no glucose). Glucose-depleted cells were grown for 1 hour at 30°C with shaking at 200rpm. After 1 hour of growth, cells were harvested by spinning in 50mL pre-chilled tubes at 3000*g*, 4°C, for 5 minutes. The supernatant was removed and the cell pellets were flash-frozen in a dry ice-ethanol bath and stored at -80°C.

### Library genomic DNA extraction

For genomic DNA extraction, between 400 million to 1.2 billion cells (two to six flash-frozen pellets) were lysed and extracted using the YeaStar Genomic DNA kit (Zymo 11-323), following the manufacturer’s instructions, with 240μL YD digestion buffer and 10μL R-Zymolyase per pellet. Extracted genomic DNA was sheared for 10 minutes (30 seconds on, 30 seconds off, on “High” setting) on ice using a Diagenode Bioruptor. Sheared gDNA was cleaned using DNA Binding Buffer (Zymo ZD4004-1-L) and UPrep Spin Columns (Genesee Scientific 88-143). Sheared and cleaned gDNA was then in vitro transcribed into RNA (denoted gRNA below and in analysis code) starting from the T7 promoter region in the insert cassette, similar to previous approaches^27,86^, using the HiScribe T7 High Yield RNA Synthesis Kit (NEB E2040S). Transcribed gRNA was cleaned using the RNA Clean and Concentrator kit (Zymo R1013).

### Library mRNA extraction

At least 200 million cells (one flash-frozen pellet) per sample was resuspended in 400μL Trizol (Thermo Fisher 15596-026) in a 1.5-ml tube and vortexed with 500μl of glass beads (Sigma G8772) at 4°C for 10 min (2 minutes on, 1 minute on ice). RNA was extracted from the resulting lysate using the Direct-zol RNA Miniprep Kit (Zymo R2070) following manufacturer’s instructions.

### mRNA and genomic DNA barcode sequencing

For pHPSC1142, pHPSC1117, and pHPSC1160 libraries, between 0.5-10μg of mRNA and gRNA for each library was reverse transcribed into cDNA using SuperScript IV (Thermo Fisher 18090050) and a primer annealing to the Illumina R1 primer binding site (oPB354). A 170 nt region surrounding the 24 nt barcode was PCR-amplified from the resulting cDNA in two rounds. Round 1 PCRs used cDNA template comprising 1/5th of the PCR reaction volume and primers oPB354 and oHP534. Round 1 PCR cycle numbers were adjusted as needed to obtain adequate product concentration while avoiding overamplification (between 5 and 15 cycles), then cleaned using DNA Binding Buffer (Zymo ZD4004-1-L) and UPrep Micro Spin Columns (Genesee Scientific 88– 343). Cleaned samples were then used as template for Round 2 PCR, and cycles were again adjusted to avoid overamplification (between 4 to 8 cycles). Round 2 PCRs used Round 1 PCR product comprising between 1/10th to 1/5th of the PCR reaction volume and oAS111 with indexed forward primers (oAS112-135 and oHP281-290). Amplified samples were run on a 2% agarose gel and fragments of the correct size were purified using ADB Agarose Dissolving Buffer (Zymo D4001-1-100) and UPrep Micro Spin Columns (Genesee Scientific 88– 343). Concentrations of gel-purified samples were measured using a Qubit dsDNA HS Assay Kit (Q32851) with a Qubit 4 Fluorometer. Samples were sequenced using an Illumina NextSeq 2000 in 1 × 50, 2 × 50, or 1 × 100 mode (depending on other samples pooled with the sequencing library). For the pHPSC1142 libraries, samples were sequenced with standard Read 1, standard Read 2, and standard i7/i5 index sequencing primers. A subset of these libraries were sent for re-sequencing to obtain greater read depth and sequenced with standard Read 1, custom Read 2 oAS1638 (to maintain compatibility with other libraries in the pool), and standard i7/i5 index sequencing primers. For the pHPSC1117 libraries, samples were sequenced with the standard Read 1 sequencing primer and standard index sequencing primers. For the pHPSC1160 libraries, samples were sequenced with standard Read 1, standard Read 2, and standard index sequencing primers.

For the FK_8_ library (pHPSC1163), between 0.5-10μg of mRNA and gRNA were reverse transcribed into cDNA using SuperScript IV and a primer annealing to the Illumina R1 primer binding site that contains a 7 nt unique molecular identifier (UMI) (oKC235). A 195 nt region surrounding the 48 nt insert was PCR-amplified from the resulting cDNA in one round using oPN776 and indexed forward primers (oKC230-233, oKC239-242). PCR cycle numbers were adjusted as needed to obtain adequate product concentration while avoiding overamplification (between 10 to 17 cycles). Amplified samples were size-selected and quantified as described previously. Samples were sequenced using an Illumina NextSeq 2000 in 1 × 70 mode using standard Read 1, custom i7 sequencing primer oKC256, standard i5, and custom Read 2 sequencing primer oKC236.

The 8× dicodon library (pHPSC1142) in glucose-depleted cells was reverse transcribed following the same procedure and primer as pHPSC1163 described above. A 219 nt region surrounding the 48 nt insert and 24 nt barcode was PCR-amplified from the resulting cDNA in one round using oPN776 and indexed forward primers (oKC230-233, oKC239-242). PCR cycle numbers were adjusted as needed to obtain adequate product concentration while avoiding overamplification (between 8 to 16 cycles). Amplified samples were size-selected and quantified as described previously. Samples were sequenced using an Illumina NextSeq 2000 in 1 × 70 mode using standard Read 1, custom i7 sequencing primer oKC256, standard i5, and custom Read 2 sequencing primer oKC236.

### Insert-barcode linkage sequencing

8–10 ng of plasmid pools (pHPSC1142, pHPSC1160, pHPSC1117) were used in PCR using Phusion polymerase (Thermo Fisher F530S) or Phusion Flash High-Fidelity PCR Master Mix (Thermo Fisher F548L). Round 1 PCR was carried out for up to 10 cycles, with 8-10 ng plasmid pool template comprising 1/5th of the PCR reaction volume, using primers oPB354 and oHP534. Round 1 PCRs were cleaned using DNA Binding Buffer (Zymo ZD4004-1-L) and UPrep Micro Spin Columns (Genesee Scientific 88–343). Cleaned samples were used as template for Round 2 PCR, carried out to between 4 to 8 cycles, using oAS111 and indexed forward primers (oAS112-135 and oHP281-290). Amplified samples were purified after size selection and quantified as described above. Samples were sequenced using an Illumina NextSeq 2000 in 2 × 50 or 1 × 100 mode. For the pHPSC1142 library, sequencing was performed using standard Read 1 sequencing primer, standard index sequencing primers, and custom Read 2 sequencing primer oAS1637. For the pHPSC1117 library, sequencing was performed using standard Read 1 sequencing primer and standard index sequencing primers. For the pHPSC1160 library, sequencing was performed using standard Read 1, standard Read 2, and standard index sequencing primers.

## Flow cytometry

Five single *S. cerevisiae* colonies integrated with plasmids described above were inoculated into separate wells of 96-well plates containing 150 μl of SCD-URA medium in each well and grown overnight at 30°C with shaking at 800rpm. The saturated cultures were diluted 100-fold into 150μl of fresh SCD-URA medium and grown for 5-6 hours at 30°C with shaking at 800rpm. The plates were placed on ice and analyzed using the 96-well attachment of a BD FACS Aria or Symphony cytometer. Forward scatter (FSC), side scatter (SSC), YFP fluorescence (FITC), and RFP fluorescence (PE.Texas.Red) were measured for 10,000 cells in each well. The resulting data in individual .fcs files for each well were combined into a single tab-delimited text file. YFP expression was first normalized to RFP expression per cell (henceforth referred to as YFP/RFP), then used to calculate the median value of each well. For the no-insert control, the median YFP/RFP values of all wells were averaged together. The median YFP/RFP value per replicate for all strains were then normalized to the average no-insert control value by taking the log2 difference. The average and standard error of this ratio across replicates were calculated (Fig. 2D).

## Computational analyses

Pre-processing steps for high-throughput sequencing were implemented as Snakemake workflows run within Singularity containers on an HPC cluster. All container images used in this study are publicly available as Docker images at https://github.com/orgs/rasilab/packages. Python (v3.9.15) and R (v4.2.2) programming languages were used for all analyses unless mentioned otherwise.

## Barcode to insert assignment

The raw data from insert-barcode linkage sequencing are in FASTQ format. All pertinent reads were concatenated into one FASTQ file using fasterq-dump --concatenate-reads, and inserts and barcodes were extracted and counted using awk (mawk implementation, v1.3.4). Only insert-barcode combinations where the insert matches a reference sequence in the list of reference sequences using awk were retained. Barcodes were aligned against themselves using bowtie2 with options -L 19 -N 1 --all --norc --no-unal -f. This self-alignment was used to exclude barcodes that are linked to different inserts or that are linked to the same barcode but are aligned against each other by bowtie2. In the latter case, the barcode with the lower count is discarded in filter_barcodes.ipynb. The final list of insert-barcode pairs is written as a comma-delimited .csv file for aligning barcodes from genomic DNA and mRNA sequencing below.

## Barcode counting in genomic DNA and mRNA

The raw data from sequencing barcodes in genomic DNA and mRNA is in FASTQ format. All pertinent reads were concatenated into one FASTQ file, and barcodes were extracted and counted using awk. For barcodes that are present in the filtered barcodes .csv file from linkage sequencing, the barcode count and associated insert are printed into a .csv file for subsequent analyses in R. For libraries containing both barcodes and UMIs, only distinct barcode-UMI combinations where the barcode is present in the filtered barcodes .csv file from linkage sequencing are retained. The number of UMIs per barcode and associated insert are printed into a .csv file for subsequent analyses in R.

## mRNA quantification and statistical analyses for barcode sequencing

Only barcodes with a minimum of 10 reads and inserts with a minimum of 2–4 barcodes were included. The mRNA level for each insert was calculated as the mean log2 ratio of the summed mRNA barcode counts to the summed gRNA barcode counts using 100 bootstrap samples. The standard deviation was calculated across all barcodes for each insert using 100 bootstrap samples. For libraries with a large number of variants (e.g. >= 70,000) mRNA levels were median-normalized within each library. For libraries with a smaller number of variants (e.g. 1000-2000), libraries were normalized to spike-in strain barcode counts or library size (RPM). For all other experiments, the standard error of the mean was calculated using the std.error function from the plotrix R package. P-values for statistically significant differences were calculated using the t.test or wilcox.test R functions as appropriate for each figure (see figure captions).

## Insert counting and mRNA quantification

For the FK_8_ deep mutational scanning library, inserts were sequenced directly and thus barcodes were not counted or used for statistical analysis. Instead, inserts and UMIs were extracted and counted using awk. Only insert-UMI combinations where the insert matches a reference sequence in the list of reference sequences using awk were retained. Subsequent insert-UMI counts were summed across the mRNA and gRNA samples. mRNA levels for each insert were calculated as the log2 ratio of the summed mRNA insert-UMI counts to the summed gRNA insert-UMI counts, and then averaged across the two biological replicates. Resultant mRNA levels were then normalized against mRNA levels of spike-in strains to allow comparison between wild-type and *hel2△* cells.

## Supplementary Figures

**Figure S1.**
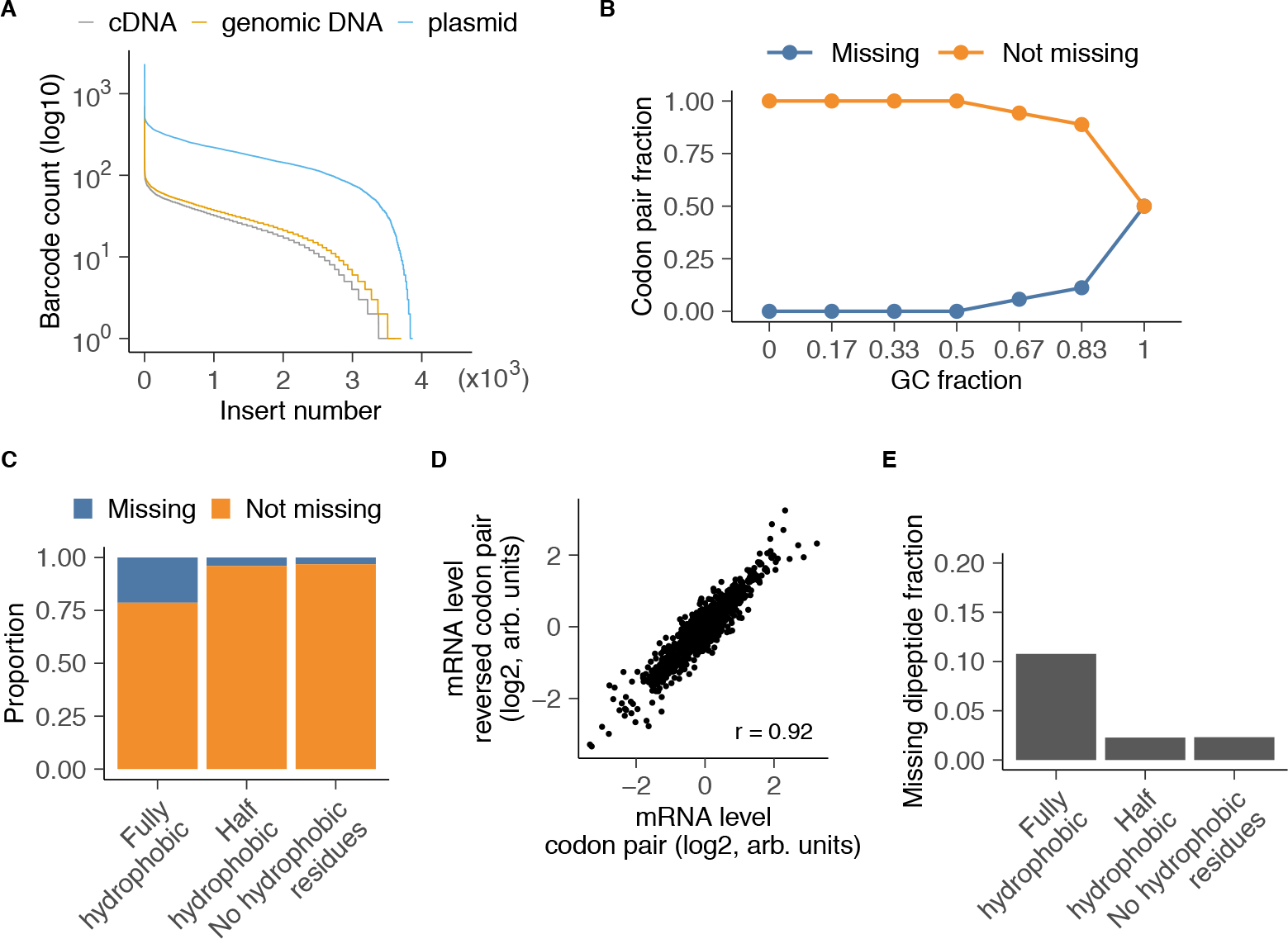
Plasmid and yeast codon pair library alignment statistics. **(A)** Distribution of barcodes per codon pair insert for the plasmid, cDNA, and genomic DNA libraries. **(B)** Proportion of missing codon pair inserts in the plasmid library by GC content. **(C)** Proportion of missing codon pair inserts in wild-type yeast grouped by hydrophobicity. **(D)** mRNA level of reporters for each codon pair compared to its reversed codon pair. Stop codon-containing pairs and pairs where the codon and reversed codon are the same are excluded. *r* indicates Pearson correlation coefficient. **(E)** Proportion of missing codon pair inserts grouped by hydrophobicity for the 139 inserts that are missing in all three strains.

**Figure S2.**
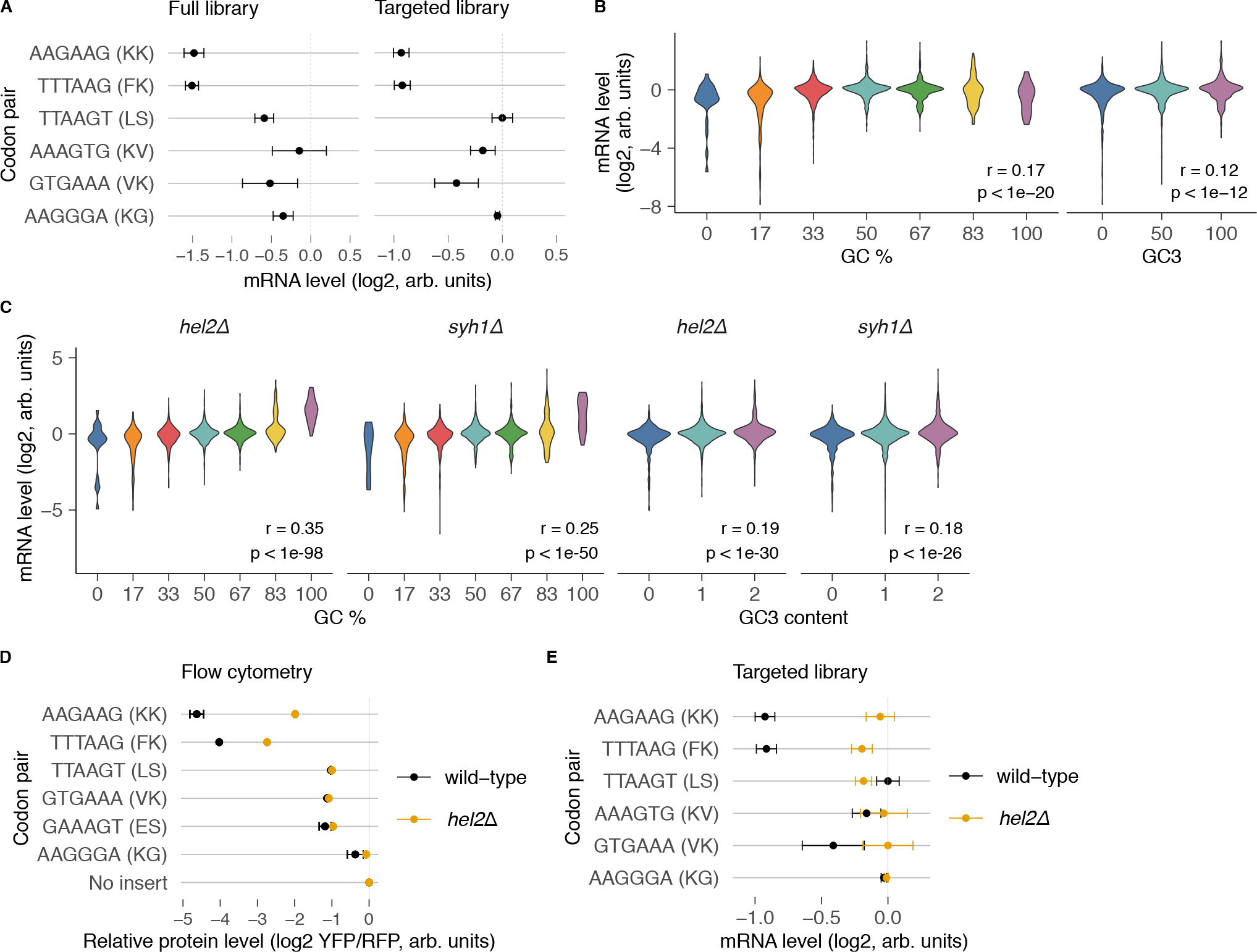
**(A)** Small-scale validation of codon pair screen. The mRNA levels for individually cloned codon pair inserts (as described in Fig. 2C, D) are plotted alongside mRNA levels for the same codon pairs taken from the full library. Error bars are calculated as in Fig. 3C. mRNA levels for the small-scale library are normalized to the maximum value and mRNA levels for the full library are normalized to the median value. **(B)** mRNA levels of codon pair inserts as a function of their GC content (left) or GC3 content (right) in wild-type cells. Pearson correlation coefficient *r* and p-value *p* are shown for GC content and GC3 content (two-sided t-test). **(C)** Same as in B, but for *hel2△* and *syh1△* cells. **(D)** Effect of individually cloned codon pair inserts on peptide expression in *hel2△* cells compared to wild-type. Peptide expression is quantified as in Fig. 2D. **(E)** mRNA level of individually cloned codon pair inserts in *hel2△* cells compared to wild-type. mRNA levels and error bars are calculated as in Fig. 3C, except with maximum-normalization.

**Figure S3.**
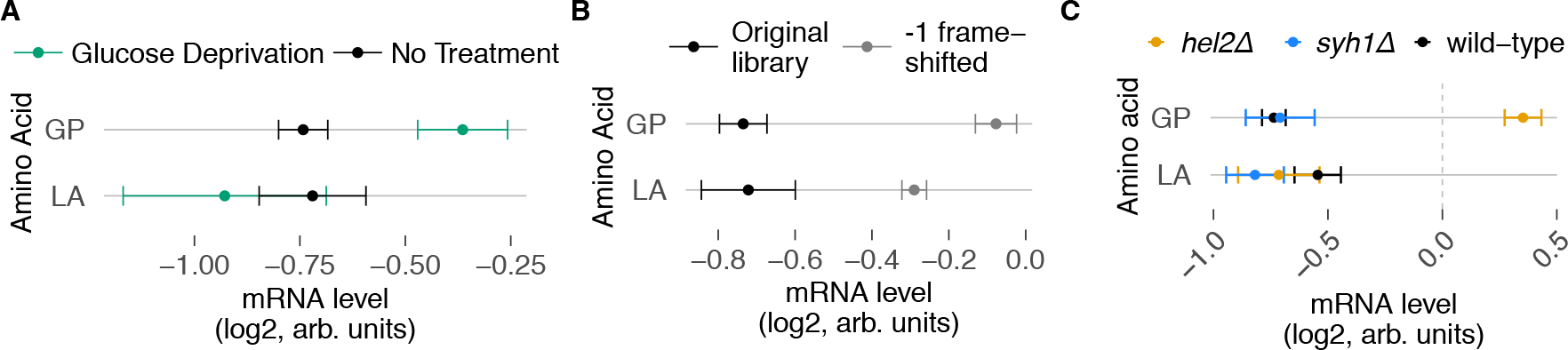
**(A)** mRNA levels for reciprocal dipeptide repeats not included in Fig. 3C. **(B)** mRNA levels for reciprocal dipeptide repeats not included in Fig. 3F. **(C)** mRNA levels for reciprocal dipeptide repeats not included in Fig. 4E.

## Supplementary Tables

**Table S1.**
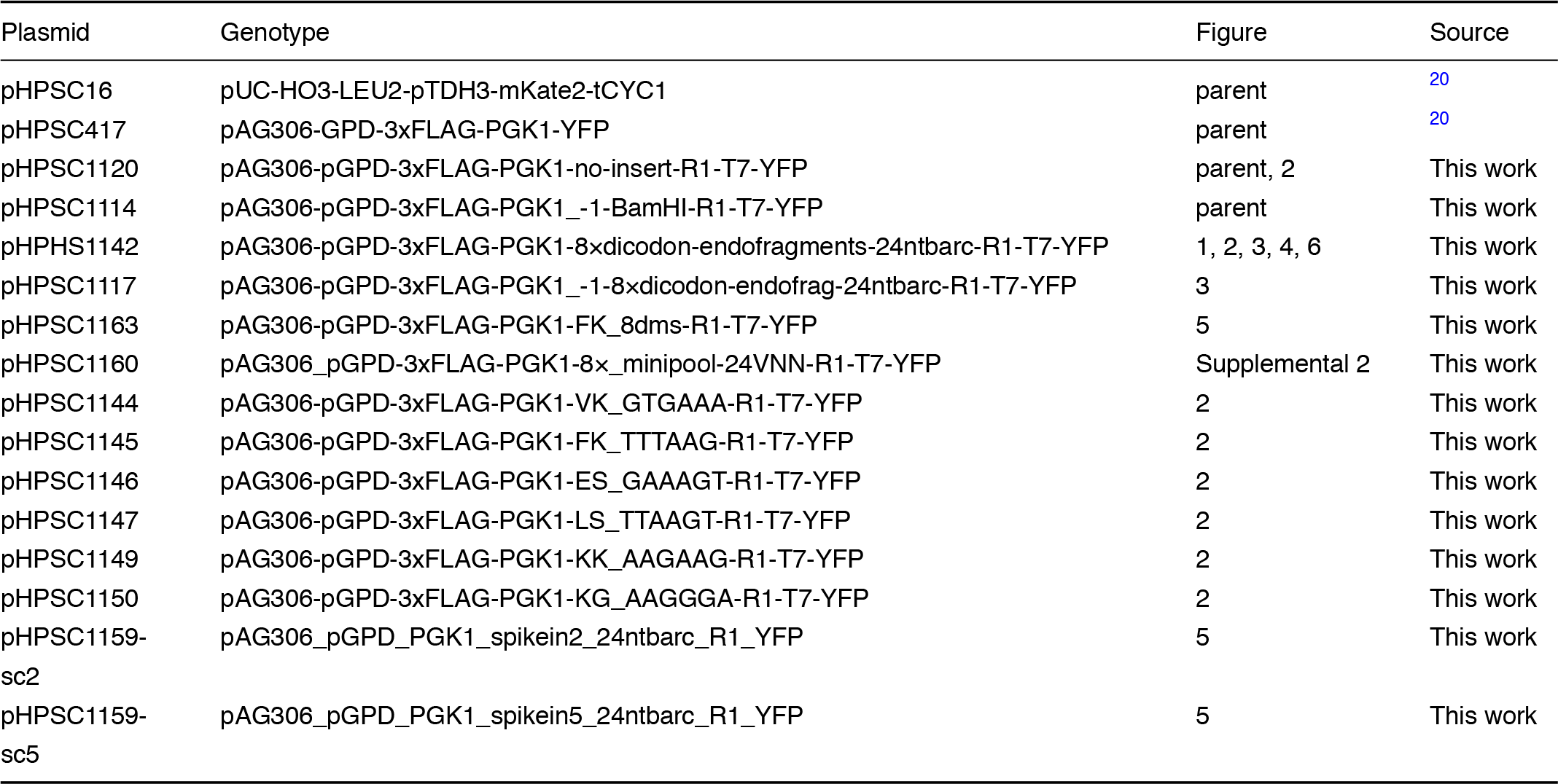
List of plasmids used for this study.

**Table S2.**
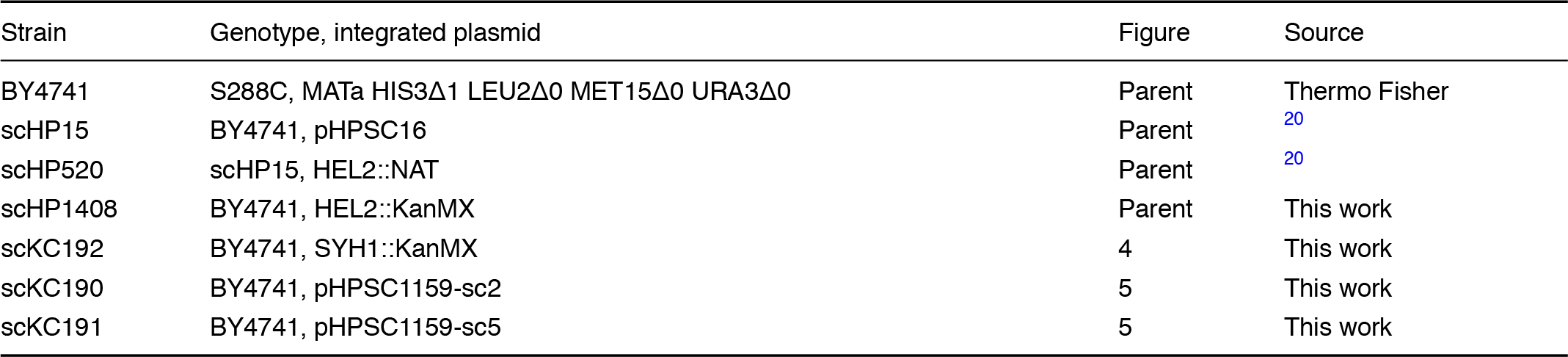
List of *S. cerevisiae* strains used for this study.

**Table S3.**
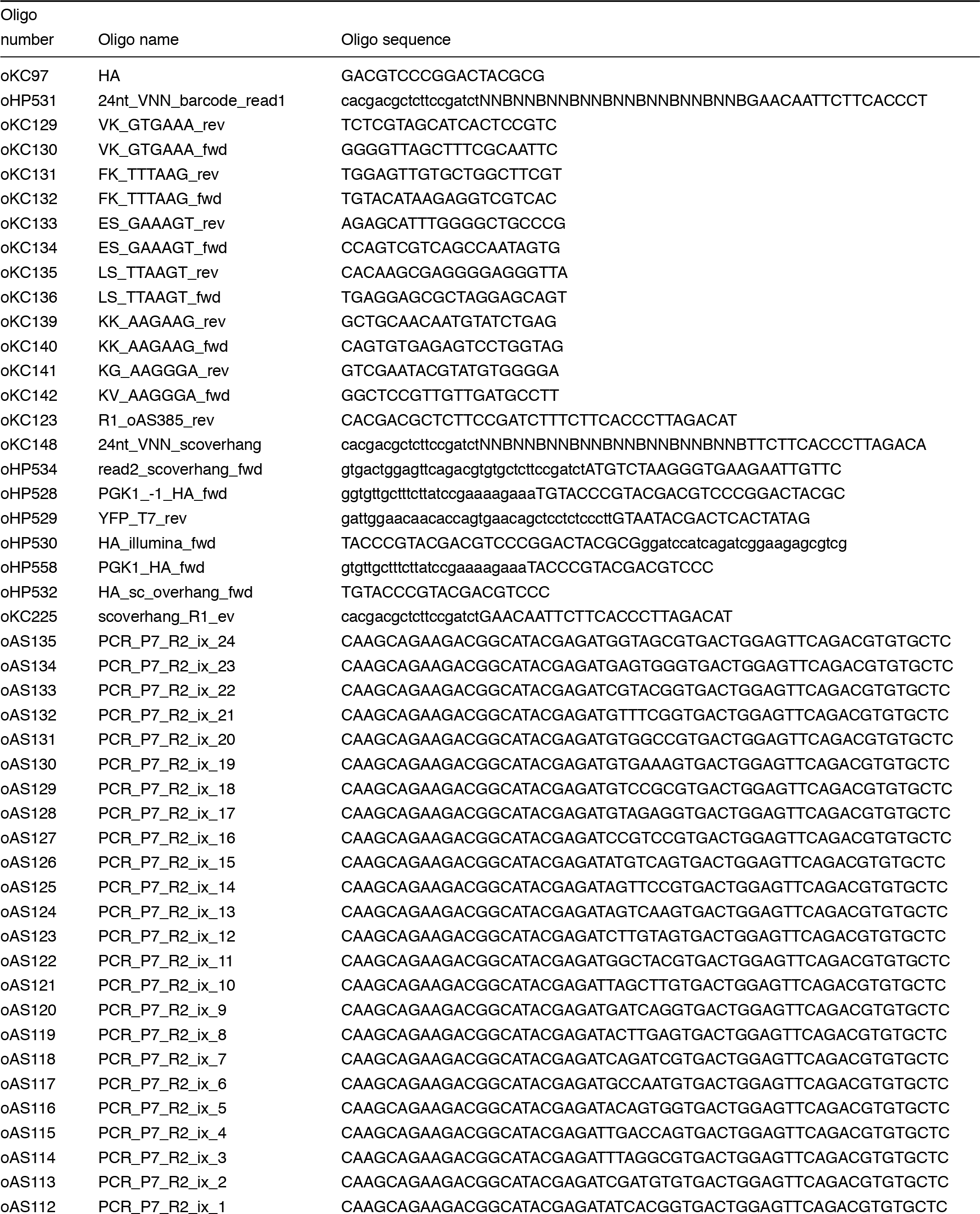
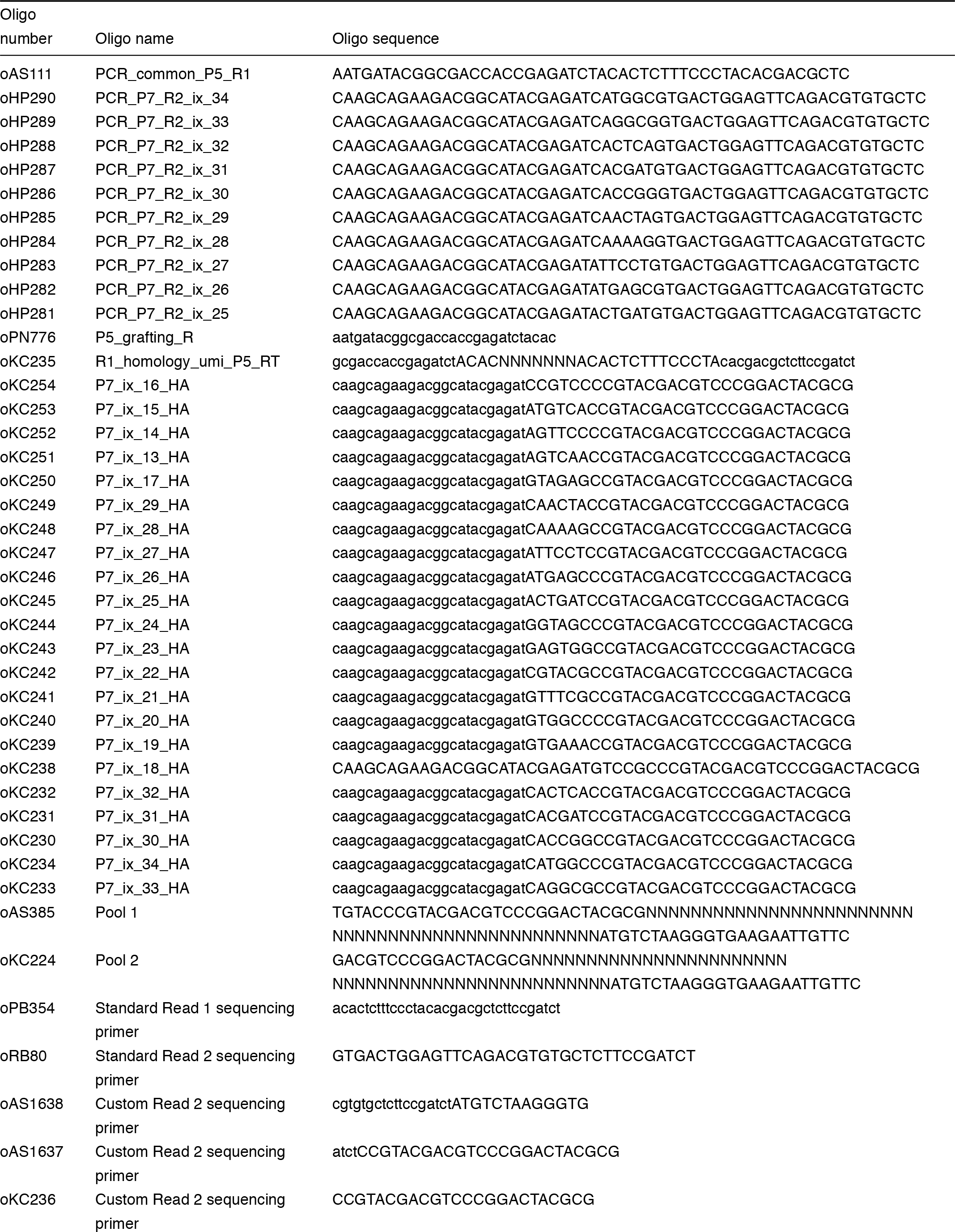
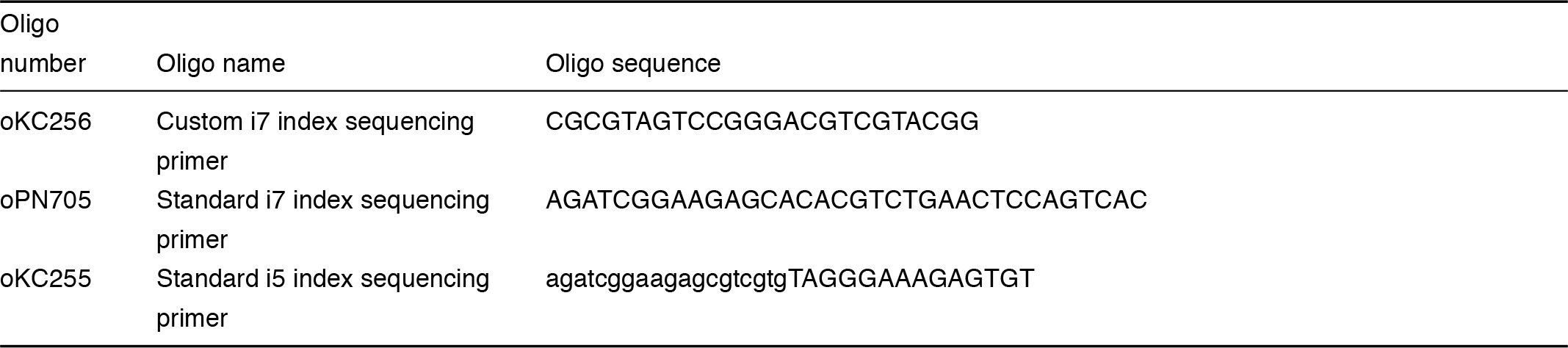
List of oligos used for this study.

